# The dysregulated IL-23/T_H_17 axis in endometriosis pathophysiology

**DOI:** 10.1101/2023.12.07.570652

**Authors:** Danielle J. Sisnett, Katherine B. Zutautas, Jessica E. Miller, Harshavardhan Lingegowda, Soo Hyun Ahn, Alison McCallion, Olga Bougie, Bruce A. Lessey, Chandrakant Tayade

## Abstract

Endometriosis is a chronic inflammatory disease where endometrial-like tissue grows ectopically, resulting in pelvic pain and infertility. Interleukin (IL)-23 is established as a key contributor in the development and differentiation of a subset of T cells known as T-helper 17 (T_H_17) cells, driving T_H_17 cells towards a pathogenic profile. In a variety of inflammatory and autoimmune disorders, such as psoriasis and rheumatoid arthritis, T_H_17 cells secrete proinflammatory cytokines including IL-17, contributing to the disease pathophysiology. Our studies and others have implicated IL-17 and T_H_17 cell dysregulation in endometriosis, which is associated with disease severity. Here we address whether IL-23 driven T_H_17 cells contribute to the cardinal features of lesion proliferation, vascularization, and inflammation in endometriosis using patient samples, representative cell lines, and our established mouse model of endometriosis. Our results indicate significantly dysregulated expression of key genes in the IL-23/T_H_17 axis in patient ectopic and eutopic endometrial samples and increased IL-23 protein in patient plasma compared to healthy controls. *In-vitro* studies using primary human T_H_ cells determined that IL-23 cocktail treatment significantly increased the frequency of pathogenic T_H_17 cells. Similarly, treatment with recombinant human (rh)IL-23 on cell lines (12Z, EECC, HUVEC, and hESC) representative of the endometriotic lesion microenvironment led to a significant increase in cytokines and growth factors known to play a role in lesion establishment and maintenance. In a syngeneic mouse model of endometriosis, treatment with recombinant mouse (rm)IL-23 led to significant alterations in numbers of myeloid and T cell subsets in peritoneal fluid and significantly increased numbers of giant cells within the lesion. Endometriotic lesions from rmIL-23 mice did not reveal significant alterations in proliferation and vascularization, although trends of increased proliferation and vascularization were observed. Collectively, these findings provide insights into the impact of the IL-23/T_H_17 axis on local immune dysfunction and broadly on the pathophysiology of endometriosis.

## Introduction

Endometriosis is characterized by the growth of endometrial-like tissue in ectopic sites and is commonly associated with dysmenorrhea, dyspareunia, chronic pelvic pain, and infertility^1,2^. While endometriosis incidence and prevalence are difficult to establish due to the invasive nature of disease diagnosis, it is estimated that endometriosis afflicts 10% (190 million) of reproductive aged women and an unknown percentage of gender-diverse individuals^3–5^. Though the exact origin and pathogenesis of endometriosis remain largely unknown, we and others postulate that immune dysfunction promotes the growth and survival of ectopic lesions by interfering with lesion clearance^1,6^. Specifically, proinflammatory IL-17 is well established to contribute to the pathogenesis of several inflammatory diseases, including endometriosis^7–10^. Indeed, we have previously shown that endometriotic lesions produce IL-17 and that patients have significantly higher IL-17 concentrations in plasma compared to controls^10^. Additionally, surgical removal of endometriotic lesions led to a significant decline in IL-17 plasma concentration, suggesting that endometriotic lesions are likely a predominant source of circulating IL-17^10^. IL-17 is largely produced by T-helper 17 (T_H_17) cells. Though, T_H_17 cells co-produce IL-17, IL-21, and IL-22, which can mediate tissue inflammation and angiogenesis^10,11^. T_H_17 cells are well documented to have a pathogenic role in various inflammatory and autoimmune disorders, such as psoriasis and rheumatoid arthritis, due to their secretion of proinflammatory cytokines which promote immune cell accumulation at inflammatory sites^11,12^. However, T_H_17 cells have a substantial degree of plasticity, adopting differing fates in response to environmental cues, such as cytokines/chemokines^8,13,14^. In fact, T_H_17 cells may also play a beneficial (non-pathogenic) role against opportunistic pathogens and promote and/or build intestinal barrier integrity^11,14–16^. Specifically, IL-17A^+^IL-10^−^ T_H_17 cells are termed “pathogenic”, while IL-17A^+^IL-10^+^ are termed “non-pathogenic”, promoting an anti-inflammatory/immunosuppressive phenotype^12,17^.

Studies show T_H_17 differentiation is initiated from naïve CD4^+^ T cells by a combination of cytokines TGF-β1, IL-1β, IL-6, IL-21, and IL-23^11,18,19^. IL-17 and IL-21 secreted by developing T_H_17 cells further drive T_H_17 cell fate and early proliferation, while IL-23 is fundamental for T_H_17 cell expansion and phenotypic stabilization^11,18–21^. Moreover, when exposed to IL-23, developing T_H_17 cells produce significantly reduced anti-inflammatory IL-10, further evoking a proinflammatory, pathogenic phenotype^17,18^. In fact, T_H_17 cells exposed to TGF-β1 and IL-6 during development are incapable of inducing autoimmune disease^18^ and unable to upregulate proinflammatory cytokines without further exposure to IL-23^17^. Hence, complete acquisition of pathogenic potential in T_H_17 cells is likely mediated by IL-23. This is supported by reports that neutralization of the IL-23-specific p19 subunit ameliorates/protects from autoimmune disease, including experimental autoimmune encephalomyelitis and autoimmune colitis, whereas loss of downstream IL-17 or IL-17RA (IL-17 receptor A) failed to reduce disease emergence/severity^11,22^. Thus, if IL-23 is a key contributor in driving T_H_17 cells to a pathogenic profile, IL-23 may govern the balance between T_H_17 and T-regulatory cells (T_REGS_) which is critical to avoid immune dysfunction and maintain immune homeostasis^23–25^.

In endometriosis, some theorize that a tolerant, anti-inflammatory peritoneal microenvironment initially allows for early endometriosis lesion development/establishment, preventing lesion clearance^26^. While later lesion growth and maintenance are supported by a proinflammatory microenvironment, promoting inflammation, angiogenesis, and fibrosis^26^. This scenario suggests a skewed T_H_17/T_REG_ axis towards a tolerant (T_REG_ dominant) microenvironment as lesions initially establish, followed by a proinflammatory (T_H_17 dominant) microenvironment allowing for lesion survival, as downstream T_H_17 cytokines, such as IL-17, are well-known to promote lesion angiogenesis, proliferation, and inflammation^10,27,28^. Results from our lab support this theory, finding T_H_17 cells to be significantly decreased in patient menstrual effluent as compared to controls^29^. Based on the theory of retrograde menstruation, this data provides insights into early events in endometriosis lesion establishment. Though further studies are warranted, this theory is also supported by findings that T_H_17 cells were significantly increased in the peritoneal fluid (PF) of patients with severe (III/IV stage) endometriosis as compared to early (I/II stage) endometriosis^13^.

IL-23 is a heterodimeric cytokine largely secreted by activated dendritic cells and macrophages^30^. IL-23 has been established to dysregulate the T_H_17/IL-17 axis in a variety of inflammatory and autoimmune disorders, such as psoriasis and rheumatoid arthritis^19,31,32^. As evidence suggests dysregulated levels of T_H_17 cells^13,24,33^ and IL-17^10,28^ in endometriosis, as well as an association of T_H_17 cells with increasing disease severity^13^, it is likely that IL-23 is regulating the T_H_17/IL-17 axis in endometriosis to exacerbate disease. While the downstream cytokine, IL-17, has been previously evaluated for its therapeutic potential and role in various autoimmune and chronic inflammatory diseases^32^, the role of upstream IL-23 has yet to be fully evaluated in endometriosis pathophysiology. In contrast to targeting IL-17, targeting IL-23 reduces proinflammatory IL-17 production by pathogenic T_H_17 cells while also preserving overall IL-17 production from other IL-17A-producing cells, ameliorating safety concerns around eliminating the protective effects of IL-17^32^. This study depicts a dysregulated IL-23/T_H_17 axis in endometriosis patient samples, which was replicated in *in-vitro* experiments. We also provide insights using a mouse model of endometriosis, revealing IL-23 to significantly alter the local immune microenvironment and T_H_17/T_REG_ axis, indicating a plausible role of IL-23 in endometriosis pathophysiology and highlighting IL-23 as a potential non-hormonal therapeutic target for patients with endometriosis.

## Methods

### Ethics statement

Ethics for the use of human samples in this study was approved by the Health Sciences Research Ethics Board at Kingston Health Sciences Centre (KHSC), Queen’s University (Kingston, ON, Canada), the Greenville Health System (Greenville, SC, USA), the University of North Carolina at Chapel Hill (Chapel Hill, NC, USA), and Wake Forest Baptist Health (Winston-Salem, NC, USA). Written, informed consent was attained from endometriosis patients and healthy controls prior to the collection and use of clinical of samples. Animal studies were approved by the Queen’s University Animal Care Committee at Queen’s University (Kingston, ON, Canada).

### Human subjects

Endometriosis patients undergoing excision surgery due to endometriosis-associated infertility and/or pelvic pain were included in this study. Healthy, fertile controls without endometriosis were recruited for endometrial biopsy and plasma collection prior to elective tubal ligations. All patients recruited for this study were free of hormonal therapy for a minimum of three months prior to sample collection.

### RNA isolation and targeted RT qPCR array for IL-23/T_H_17/IL-17 axis-associated genes in patient tissues

Total RNA was extracted from ectopic (n=9) and eutopic (n=8) endometrium of patients, and normal healthy endometrium (n=9) using a total RNA purification kit (17200, Norgen Biotek Corp., Canada) as per manufacturer’s protocol. RNA quality was assessed and cDNA synthesized as previously detailed^34^. RT qPCR was conducted with custom array plates (330171, Qiagen, Canada) using the LightCycler 480 Real-Time PCR system (Roche Molecular Systems Inc., Switzerland) with RT^2^ SYBR Green qPCR Mastermix (330501, Qiagen, Canada) and 200ng cDNA, as per manufacturer’s protocol. Data analysis was conducted using the ΔΔCt method and relative quantification was performed by geometric averaging of *Gapdh* and *Actb* reference genes. The custom RT qPCR array was designed to target 11 genes of interest in the IL-23/T_H_17/IL-17 axis: *IL23A, IL12B, IL12A, IL23R, IL21, IL22, IL17A, IL17F, IL17RA, IL6, TGFB1*.

### ELISA for detection of IL-23 in patient tissue and plasma samples

IL-23 concentration was measured in endometriosis patient tissue protein extracts (ectopic [n=10] and eutopic [n=16] endometrium) and patient plasma (n=13) as compared to healthy controls (n=19), as well as between stratified patient plasma samples (n=7) over three time points using an IL-23 Human ELISA Kit (BMS2023-3, ThermoFisher, Canada), as per manufacturer’s protocol. 20mg of tissues were homogenized in PowerBead tubes (13113-50, Qiagen, Canada) containing protease inhibitor cocktail (535140, Sigma-Aldrich, Canada) and tissue protein extraction reagent (78510, ThermoFisher, Canada). Protein concentration was determined using a Pierce™ BCA protein assay kit (23225, ThermoFisher, Canada) as per kit protocol. Samples were normalized to the lowest concentration and stored at -80°C prior to analysis. ELISA results were read in a SpectraMax iD5 microplate reader (Molecular devices, USA) at an absorbance of 450nm and a reference wavelength of 620nm.

### Immunohistochemistry for IL-23 in an endometrioma tissue microarray

An endometrioma tissue microarray (TMA) was constructed using patient tissue samples from Kingston General Hospital (Kingston, ON, Canada), as previously described^35^. Matched patient ectopic and eutopic tissues (n=17) as well as normal endometrium controls (n=10) were subjected to immunohistochemistry (IHC) staining using a polyclonal IL-23 antibody (1:200; ab45420, Abcam, UK) and analysis using HALO image analysis software (Indica Labs, USA) was conducted as previously outlined^34^.

### Primary human CD4^+^ T cell isolation, *in-vitro* T_H_17 polarization, and multiplex cytokine array

30mL of whole blood was collected from healthy volunteers into BD Vacutainer® tubes (CABDL366643L, VWR, USA). Peripheral blood mononuclear cells (PBMCs) were isolated using Lymphoprep™ (07811, StemCell Technologies, Canada) and SeptMate™-50 PBMC Isolation Tubes (85460, StemCell Technologies, Canada). Primary CD4^+^ T cells were isolated from PBMCs using an EasySep Human CD4^+^ T Cell Isolation Kit (17952, StemCell Technologies, Canada) as per manufacturer’s instruction. Primary CD4^+^ T cells were seeded at 1×10^6^ cells/well in 24-well plates and maintained in X-VIVO-15™ media (CA12001-988, VWR, USA). Four treatment groups were used: i) cocktail and activation, ii) activation alone, iii) cocktail alone, and iv) media control.

Activation consisted of 10μg/mL plate-bound anti-human CD3 antibody (300437) and 1μg/mL soluble anti-human CD28 antibody (302902) purchased from BioLegend, USA. Based on previously published protocols^36–39^, the T_H_17 cocktail consisted of 5ng/mL recombinant human TGF-β1 (rhTGF-β1; 240-B-002), 12.5ng/mL rhIL-1β (201-LB-005), 25ng/mL rhIL-6 (206-IL-010), 25ng/mL rhIL-21 (8879-IL-010), 25ng/mL rhIL-23 (1290-IL-010), 500ng/mL anti-IFNγ (506531, BioLegend, USA), and 500ng/mL anti-IL-4 (500837, BioLegend, USA) purchased from R&D Systems unless stated otherwise. Primary CD4^+^ T cells were treated in triplicates and incubated at 37°C with 5% CO_2_ for 4 days, as per established protocols^36,37^. On day 4, cells were re-stimulated with 50ng/mL of PMA (74042, StemCell Technologies, Canada) and 250ng/mL of ionomycin (73722, StemCell Technologies, Canada) in combination with 2μL/mL of protein transport inhibitor cocktail (Brefeldin A and Monensin; 00-4980-93, ThermoFisher, Canada) and incubated at 37°C with 5% CO_2_ for 5hrs prior to analysis. Supernatant was analyzed using a commercially available multiplex cytokine/chemokine analysis (HD48-plex, EveTechnologies, Canada). To examine cell fate following *in-vitro* polarization, flow cytometry was conducted on day 4.

### Human cell lines

An immortalized human endometriotic epithelial cell line derived from epithelial cells of peritoneal endometriosis (12Z; kindly gifted by Dr. Anna StarzinskiPowitz), endometrial epithelial carcinoma cells (EECC; ATCC CRL-1671), human umbilical vein endothelial cells (HUVEC; ATCC CRL-1730), and human endometrial stromal cells (hESCs; T0533 ABM, Canada) were utilized in this study. EECCs were maintained in DMEM (D6429, Sigma-Aldrich, Canada) supplemented with 10% FBS (FBS; 97068-085, VWR, USA) and 1% penicillin and streptomycin (15140122, ThermoFisher, Canada). Cell line cultivation as well as 12Z, HUVEC, and hESC maintenance was conducted as previously described^34^.

### Multiplex cytokine analysis of rhIL-23 treated endometriosis representative cell lines

12Zs, EECCs, HUVECs, and hESCs were seeded at 3×10^5^ cells/well onto 6-well plates and rested for 24hrs prior to stimulation with PBS or rhIL-23 (1290-IL-010, R&D Systems, USA) at concentrations of 1, 10, 50, and 100ng/mL. Cells were treated in triplicates and incubated at 37°C with 5% CO_2_ for 24hrs and supernatant was analyzed via multiplex cytokine array (HD48-plex, EveTechnologies, Canada).

### Proliferation and apoptosis assays in rhIL-23 treated human cell lines

12Zs, EECCs, HUVECs, and hESCs were seeded onto 96-well plates at 5×10^3^ cells/well and rested for 24hrs. Cells were then treated in triplicates for 24hrs with varying concentrations of rhIL-23 for analysis via a WST-1 proliferation assay using WST-1 reagent (0501594400, Sigma-Aldrich, Canada) and an apoptosis assay using Caspase-Glo® 3/7 reagent (G8091, Promega, Canada), as per manufacturer’s protocol. Results of WST-1 and Caspase-Glo® assays were acquired as previously outlined^34^.

### Endothelial cell tube formation assay

To determine the influence of IL-23 on angiogenesis, a tubulogenesis assay was conducted using IBIDI μ-Plates (81506, IBIDI, Germany), as per manufacturer’s protocol. Briefly, each μ-Plate well was coated with 10 μL of Matrigel™ (353230, Corning, USA) and incubated for 1hr in a humid chamber at 37°C with 5% CO_2_. HUVEC cells were then seeded at 1×10^4^ cells/well in triplicates on top of polymerized Matrigel™ in 50uL of media containing either VEGF (25ng/mL), PBS (vehicle), or rhIL-23 (1, 10, 50, 100ng/mL). Cells were incubated for 4hrs at 37°C with 5% CO_2_ prior to image acquisition. Images were captured at 4X objective using an Olympus CKX41 Microscope (Olympus Life Science, USA) with a Lumenera Infinity 1-3 microscope camera. Images were analyzed for tube formation using WIMASIS Software (WimTube: Tube Formation Assay Image Analysis Solution)^40^.

### Murine *in-vitro* T_H_17 polarization

Spleens from naïve female C57BL/6 mice (n=6) were collected and mechanically digested through 70μm strainers before centrifugation at 400RCF for 5mins at 4°C and pellets were resuspended and counted. An EasySep™ Isolation Kit (19765, StemCell Technologies, Canada) was used to isolate naïve CD4^+^ T cells, following manufacturer’s protocol. CD4^+^ T cells were washed with 2% FACs buffer (PBS with 2% FBS) prior to resuspension in TexMACS complete media (130-097-196, Miltenyi Biotec, USA), supplemented with 10% FBS, 1% penicillin and streptomycin, and 0.1mM 2-ME (Mercaptoethanol; 31350-010, ThermoFisher, Canada). Cells were counted and seeded at 2.5×10^5^ cells/well in 96-well plates. Four treatment groups were used: i) cocktail and activation, ii) cocktail and activation supplemented with an additional 10ng/mL recombinant mouse (rm)IL-23, iii) 10ng/mL rmIL-23 (no cocktail) and activation, and iv) media and activation (control). Activation consisted of 5μg/mL plate-bound anti-mouse CD3ε antibody (100340) and 5μg/mL soluble anti-mouse CD28 antibody (102116) purchased from BioLegend, USA. A commercially available CytoBox T_H_17 mouse cocktail (130-107-758, Miltenyi Biotec, USA) was used for polarization of murine CD4^+^ T cells *in-vitro*, as per manufacturer’s instruction. To examine cell fate, flow cytometry was conducted on day 5 following re-stimulation for 5hrs with 50ng/mL PMA, 750ng/mL ionomycin, and 2μL/mL protein transport inhibitor cocktail (last 4hrs).

### Murine model of endometriosis

Endometriosis was induced in seven-to-eight-week-old female C57BL/6 mice (n=12; Charles River Laboratories, USA) and sham surgery was conducted (n=12) as previously detailed^34^. Post-surgical care was conducted as outlined^28^. To examine the effects of IL-23 *in-vivo*, both endometriosis and sham mice were treated with intraperitoneal (i.p.) injections of 1μg of rmIL-23 (n=6/group; 589006, BioLegend, USA), or PBS for controls (n=6/group), three times a week for three weeks (**Figure 6A**). Mice were euthanized 21 days post-op by 5% isoflurane inhalation followed by cervical dislocation. PF, endometriosis-like lesions, uterine horns, and spleens, were collected, processed, and stored prior to analysis as previously detailed^34^. For flow cytometry, PF and splenocytes were thawed and resuspended in 10% FACs. Cells were centrifuged at 300RCF for 5mins at 4°C and supernatant was decanted before resuspension in 100μg/mL DNAse I (10104159001, Sigma-Aldrich, Canada), as per manufacturer’s protocol. Cells were neutralized with 10% FACs, centrifuged at 300RCF for 5mins at 4°C, and supernatant was decanted before resuspension in 2% FACs for flow cytometry.

### Immunohistochemistry and H&E staining of murine lesions

Paraformaldehyde-fixed, paraffin-embedded tissue blocks were sectioned at 4μm thickness and deparaffinized using citrasolv and rehydrated in graded concentrations of ethanol. Following antigen retrieval for 20mins using citrate buffer and blocking of endogenous peroxidase activity, sections were stained with polyclonal antibodies for anti-mouse Ki67 (1:1500; ab15580, Abcam, UK) and anti-mouse CD31 (1:100; 77699S, New England Biolab, USA) to assess proliferation and vascularization, respectively, and digitally scanned for analysis as previously described^34^. A computer-generated algorithm for Ki67 and CD31 was used to digitally analyze lesions. Custom trained Cytonuclear and Area Quantification modules were used to quantify Ki67 and CD31 staining, respectively. Standard H&E staining was conducted as previously explained^41^.

### *In-vivo* mouse model to prime T_H_17 cell derivation/polarization

IL-6 and TGF-β1 are known to be key players in the initial derivation of murine T_H_17 cells, while IL-23 is vital for the establishment and clonal expansion of pathogenic of T_H_17 cells^39,42^. Thus, to assess whether T_H_17 cells may be primed *in-vivo* during lesion development, endometriosis was surgically induced in recipient mice (n=16) as described^34^. Endometriosis-induced mice either received daily i.p. injections of a T_H_17 priming cocktail (n=8; 0.5μg rmIL-6 [575704, BioLegend, USA] and 1μg rmTGF-β1 [763104, BioLegend, USA]) or rest (n=8) from days 0-3 prior to daily i.p. injections of 1μg rmIL-23 (n=4/group) or PBS (n=4/group) for 10 days (**Figure 8A**). Mice were euthanized 14 days post-op and PF, spleen, and endometriosis-like lesions were collected. Lesions were snap-frozen in liquid nitrogen prior to storage at -80°C for later analysis and PF was processed for cryopreservation, as detailed^34^, for later analysis via flow cytometry. Splenocytes were isolated as previously outlined^34^ prior to subjection to ACK lysis buffer (A1049201, ThermoFisher, Canada), per manufacturer’s instruction. Splenocytes were neutralized with RPMI-1640 (11875085, ThermoFisher, Canada) and centrifuged at 400RCF for 5mins at 4°C prior to resuspension and counting. Samples were seeded in 24-well plates at 1×10^6^ cells/well and incubated at 37°C with 5% CO_2_ with 50ng/mL PMA and 500ng/mL ionomycin for 4hrs to stimulate intracellular cytokine production. After 1hr of incubation, 2μL/mL of protein transport inhibitor cocktail was added. Flow cytometry was conducted to examine cell fate.

### Flow cytometric analyses

Cell suspensions were counted and aliquoted (5×10^5^ cells/sample) before staining with Human TruStain FcX™ (422302) or TruStain FcX™ (101319) to block non-specific antigen binding, as per manufacturer’s instruction. All products were purchased from BioLegend, USA unless otherwise stated. Cells were stained with extracellular antibodies as detailed in **Table 1**. To assess viability, human samples were stained with eBioscience™ Fixability Viability Dye eFluor™ 780 (65-0865-18, ThermoFisher, Canada), while murine samples were stained with Zombie Green™ Fixable Viability Dye (423111). All cells were then fixed and permeabilized using the eBioscience™ FoxP3/Transcription Factor Staining Buffer Set (00-5523-00, ThermoFisher), as per manufacturer’s instruction. Following permeabilization, cells were stained with intracellular antibodies (**Table 1**).

**Table 1:** List of extracellular and intracellular antibodies used for flow cytometric analyses.

| Antibody | Fluorophore(s) | Verified Reactivity | Clone | Catalogue Number | Vendor |
| --- | --- | --- | --- | --- | --- |
| CD45 | PB;<br>APC/Cy7 | Human;<br>Mouse | HI30;<br>30-F11 | 304029;<br>103115 | BioLegend, USA |
| CD3 | Alexa Fluor® 700 | Mouse | 17A2 | 100215 | BioLegend, USA |
| CD4 | PE/Cy7 | Human;<br>Mouse | RPA-T4;<br>GK1.5 | 300512;<br>100421 | BioLegend, USA |
| CD11b | Alexa Fluor® 700 | Mouse, Human,<br>Cynomolgus,<br>Rhesus monkey | M1/70 | 101222 | BioLegend, USA |
| CD25 | Brilliant Violet™ 650 | Mouse | PC61 | 102038 | BioLegend, USA |
| CD196<br>(CCR6) | FITC;<br>PE/Dazzle™ 594 | Human,<br>Cynomolgus,<br>Rhesus monkey;<br>Mouse | G034E3;<br>29-2L17 | 353412;<br>129822 | BioLegend, USA |
| ROR-γt | APC | Human, Mouse,<br>Rhesus monkey | AFKJS-9 | 17-6988-82 | ThermoFisher,<br>Canada |
| FoxP3 | PE | Mouse | MF-14 | 126404 | BioLegend, USA |
| IL-17A | PE;<br>Brilliant Violet™ 421 | Human;<br>Mouse | BL168;<br>TC11-18H10.1 | 512306;<br>506926 | BioLegend, USA |
| IL-10 | Brilliant Violet™ 605 | Mouse | JES5-16E3 | 505031 | BioLegend, USA |
| IFNγ | PE | Mouse | XMG1.2 | 505807 | BioLegend, USA |

Cells were washed with 2% FACs and data was acquired using CytoFLEX S (Beckman Coulter, USA) and analyzed via FlowJo v10 software (Oregon, USA). FMO controls were used appropriately to determine gating strategy. To set a viability gate, an aliquot of heat-killed cells was combined 1:1 with live cells and stained with viability dye.

### Statistics

GraphPad Prism9®, USA was used for all statistical analyses. When appropriate, an unpaired, non-parametric student’s t-test with Mann-Whitney correction was used to analyze two groups and a one-way ANOVA with Tukey post-hoc was used to compare 3 or more groups. Error bars signify the standard deviation (SD), and statistical significance was denoted by a p-value ≤ 0.05.

## Results

### The IL-23/T_H_17 axis is dysregulated in endometriosis patient tissues and plasma

To our knowledge, this is the first report quantifying RNA and protein expression of IL-23 [p19 (*IL23A*) and p40 (*IL12B*) subunits] and key mediators in the IL-23/T_H_17 axis in endometriosis patient ectopic (lesion) and matched eutopic endometrial tissues as compared to healthy controls. We first aimed to determine if endometriotic lesions are a source of IL-23 and how gene expression differs between matched patient ectopic and eutopic tissues. To do so, a RT qPCR array was designed to target 11 key genes of interest known to play a role in the IL-23/T_H_17 axis (**Figure 1**). We discovered that *IL23R, IL17A, IL17F, IL21, IL22, IL12B*, and *TGFB1* were significantly upregulated in patient ectopic tissues as compared to both matched eutopic tissues and healthy controls (**Fig. 1A-B**). Notably, *IL23R* (IL-23 receptor) was also significantly upregulated in patient eutopic tissues as compared to controls (**Fig. 1C**).

**Figure 1:**
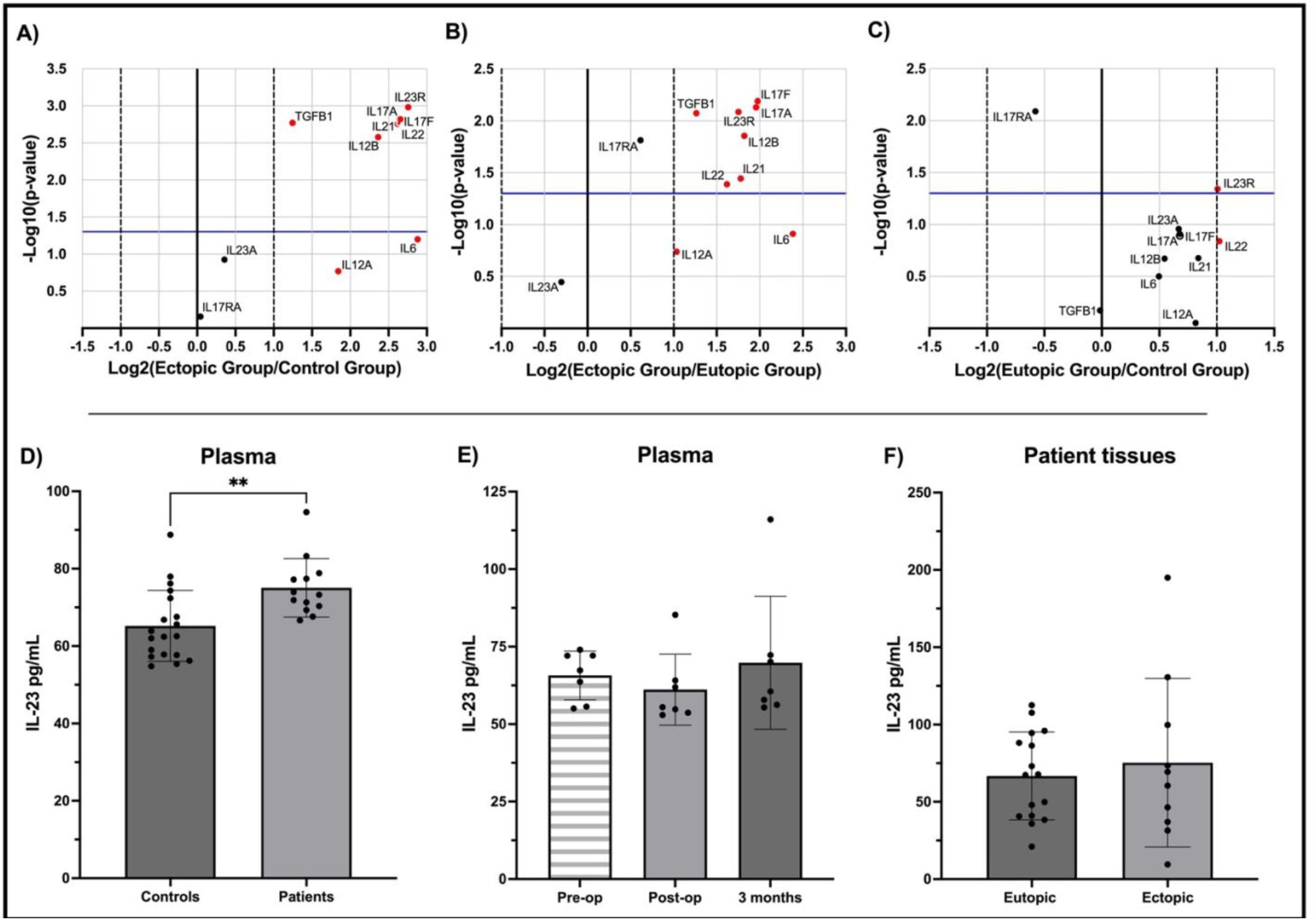
Endometriosis patient tissue samples reveal significantly dysregulated expression of key genes in the IL-23/T_H_17 axis and IL-23 protein was significantly increased in the plasma of patients as compared to healthy controls. Volcano plots depict differential mRNA expression of key genes in the IL-23/T_H_17 axis within patient ectopic tissues (n=9) vs controls (n=9; **A**), patient ectopic vs eutopic tissues (n=8; **B**), and patient eutopic vs control endometrium (**C**). The horizontal blue line indicates a significance of p=0.05 and the dashed vertical lines indicate a fold change of ± 1, whereby upregulated genes are displayed in red. IL-23 protein was measured in patient plasma, and tissues (eutopic and ectopic) as compared to healthy controls via ELISA. IL-23 concentration was significantly increased in endometriosis patient plasma (n=13) as compared to controls (n=19; **D**). However, no significant differences were found when stratifying patient plasma (n=7) by timepoints (**E**). IL-23 protein was also detected in patient eutopic (n=16) and ectopic tissues (n=10), though no significant differences were found (**F**). Data is represented as mean ± SD. Statistical analyses were conducted by a non-parametric student’s t-test with Mann-Whitney correction (D), a paired, non-parametric, one-way ANOVA with Friedman post-hoc (E), or a paired, non-parametric Student’s t-test (Wilcoxon test; F). **p<0.01.

Using ELISA, we found that IL-23 protein was significantly increased in endometriosis patient plasma samples compared to controls (**Fig. 1D**). Patient plasma was then stratified by timepoints, specifically pre-operative, post-operative, and 3 months following endometriosis lesion excision surgery to determine if lesion presence alters circulating IL-23 levels. Though, no significant differences were found between timepoints (**Fig. 1E**). IL-23 protein was also detected in patient ectopic lesion and eutopic samples, though no significant differences were revealed when comparing eutopic and ectopic tissues (**Fig. 1F**).

We then aimed to depict spatial localization of IL-23 within a human TMA (**Figure 2**) containing matched patient ectopic (endometrioma) and eutopic tissue samples (n=17; **Fig. 2A-B**) as compared to healthy control endometrium (n=10; **Fig. 2C**). IL-23 protein expression was quantified using area quantification of percent positive stain of anti-IL-23 stain across total core area (**Fig. 2D**) and independently within epithelial (luminal and glandular) and stromal tissue (**Fig. 2E**). While there were no significant differences in IL-23 protein expression between matched patient (ectopic/eutopic) or control endometrium, IHC staining illustrates that IL-23 is predominately expressed in epithelial tissues as compared to stroma of matched patient (ectopic/eutopic) and control endometrium.

**Figure 2:**
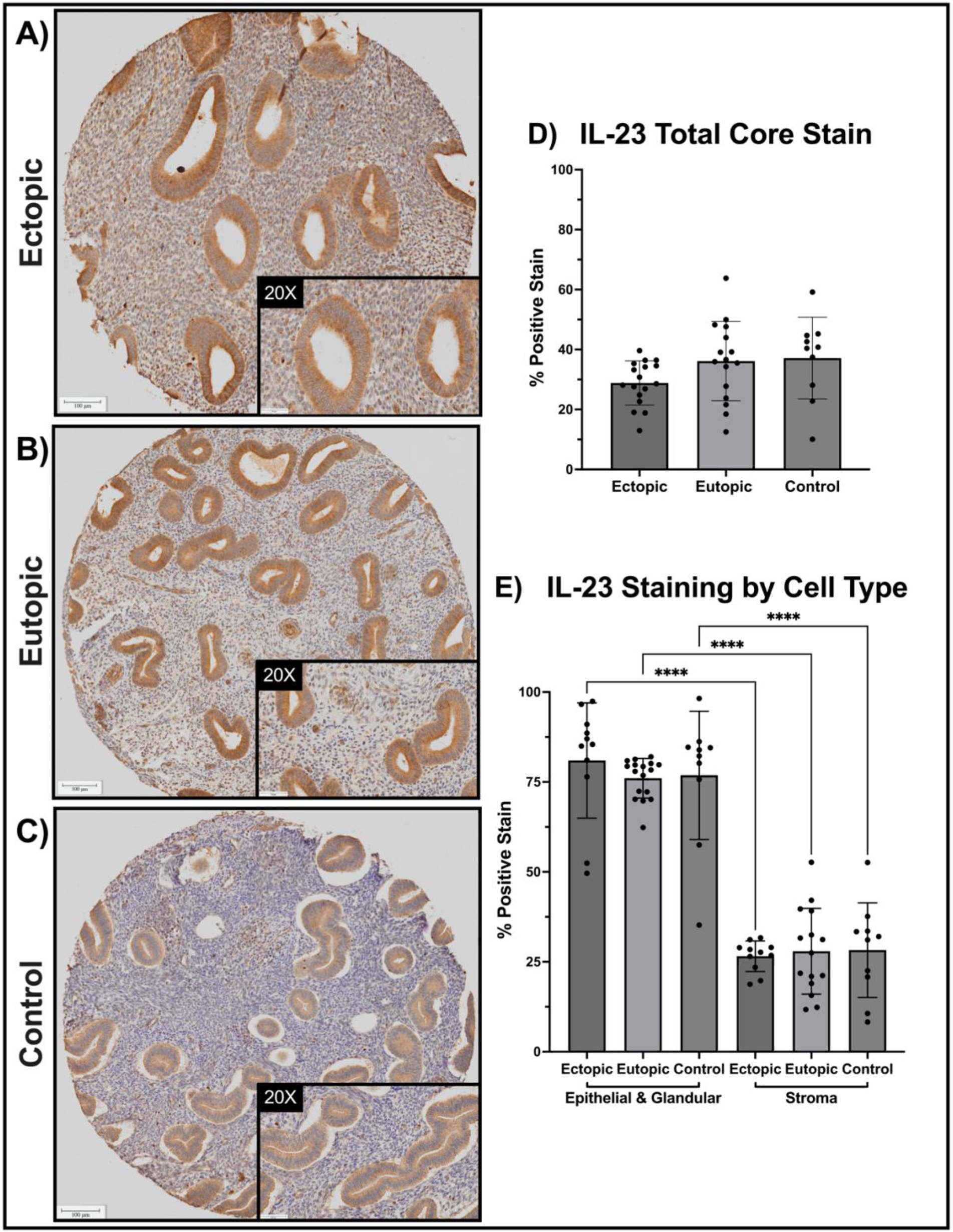
IL-23 protein is expressed in endometriosis lesions and is significantly increased in luminal and glandular epithelium as compared to stroma. Anti-IL-23 stained endometrioma tissue microarray of matched endometriosis (eutopic and ectopic, n=17; **A-B**) and control endometrium (n=10; **C**). Area quantification of percent positive stain was calculated for total core area (**D**) and luminal and glandular epithelium as compared to stromal tissue **(E**). Statistical significance was assessed using a one-way ANOVA with Tukey post-hoc. ****p<0.0001. Scanned IHC images were digitally analyzed using HALO Imaging Software (Indica Labs, USA) and provided at 5x and 20x magnification; scale bars 100μm and 50μm respectively.

### IL-23 cocktail treatment significantly alters the production of pathogenic T_H_17 cells in humans and IL-23 alone increases factors known to play a role in lesion establishment and maintenance

In accordance with literature, we sought to investigate the involvement of IL-23 in the derivation of human pathogenic T_H_17 cells using flow cytometry (**Figure 3A**) and multiplex cytokine array. We found primary human CD4^+^ T cells treated *in-vitro* with IL-23 cocktail (rhIL-23, rhIL-21, rhTGF-β1, rhIL-1β, rhIL-6, anti-IFNγ, and anti-IL-4) produced a significantly increased frequency and number of T_H_17 cells (**Fig. 3B-C**). We then aimed to determine how T_H_17 cells driven in the presence of IL-23 may influence the endometriotic microenvironment. Results from the multiplex cytokine array revealed that when activated and driven in the presence of IL-23, T_H_17 cells produced significantly increased IL-17A/IL-17F production and significantly decreased IL-10 production, suggesting a pathogenic phenotype (**Fig. 3D-E & Supplemental Fig. 1A**). Activated T_H_17 cells driven in the presence of IL-23 also secreted significantly increased monocyte chemoattractant protein-1 (MCP-1), chemokine (C-C motif) ligand 5 (CCL5/RANTES), chemokine (C-X-C motif) ligand 9 (CXCL9), and macrophage-derived chemokine (MDC) (**Fig. 3F-G & Supplemental Fig. 1B-C**).

**Figure 3:**
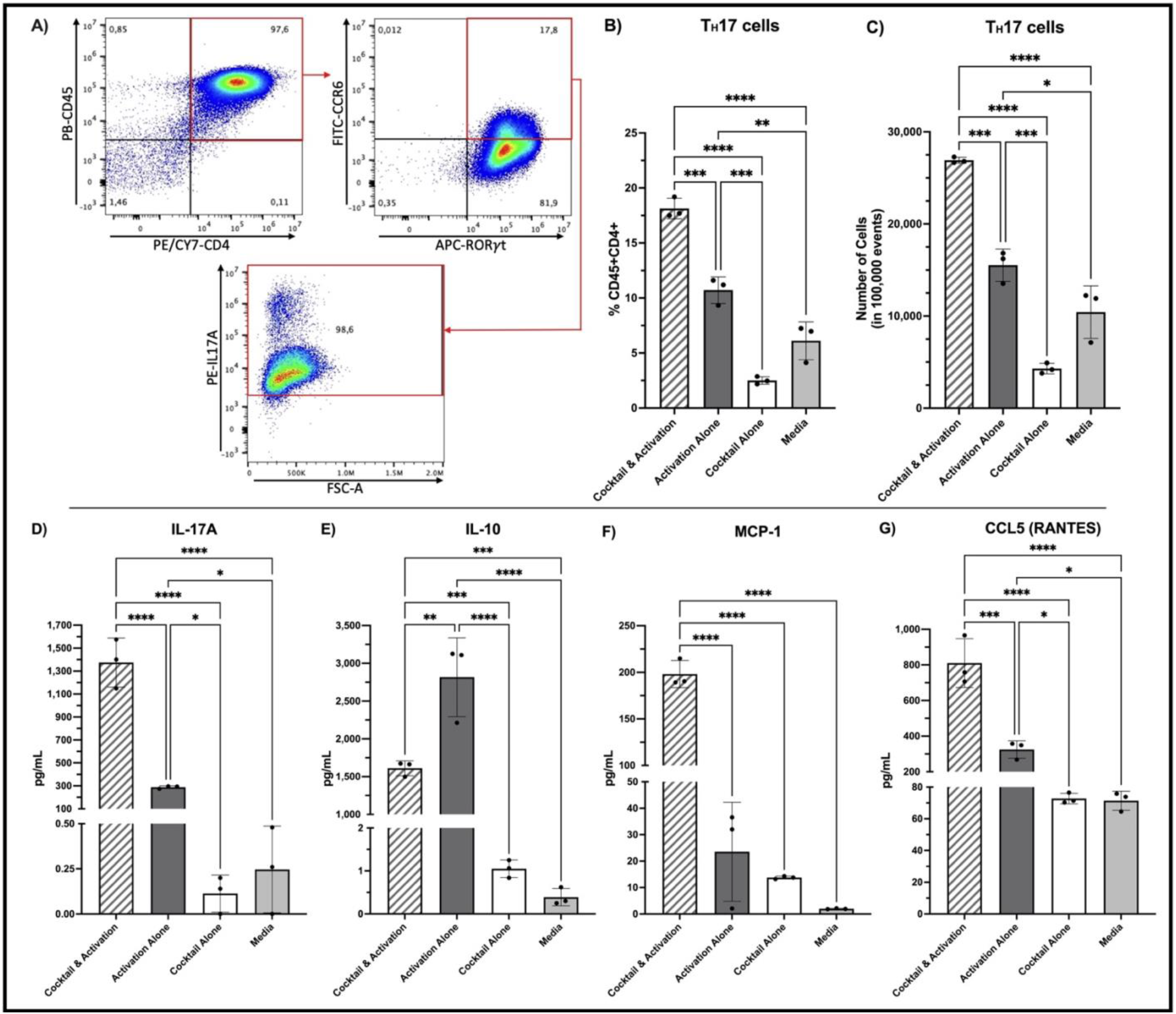
IL-23 cocktail treatment significantly promotes the production of pathogenic T_H_17 cells from primary human naïve CD4^+^ T cells and alters the production of factors involved in endometriosis lesion establishment and maintenance. Primary naïve CD4^+^ T cells isolated from human PBMCs were treated in-vitro in triplicates for 4 days with an IL-23 cocktail and CD3/CD28 activation, CD3/CD28 activation alone, IL-23 cocktail alone, or media. Cells were re-stimulated with PMA/ionomycin and protein transport inhibitor cocktail prior to supernatant collection and flow cytometric analysis for T_H_17 cells (CD45^+^CD4^+^CCR6^+^RORγt^+^IL17A^+^; **A**). Primary CD4^+^ T cells treated with cocktail and activation produced a significantly higher frequency of T_H_17 cells within the CD45^+^CD4^+^ cell population (**B**) and number of T_H_17 cells per 100,000 events (**C**), as compared to all other treatment groups. Primary CD4^+^ T cells treated with cocktail and activation were confirmed to be acting as pathogenic T_H_17 cells via multiplex cytokine array. All 48 analytes measured in supernatant showed significant differences, including IL-17A, IL-10, MCP-1, and CCL5 (RANTES; **D-G**). Data is represented as mean ± SD. A one-way ANOVA with Tukey post-hoc was used to assess significance. *p<0.05, **p<0.01, ***p<0.001, ****p<0.0001.

We then sought to determine the independent effects of IL-23 on cell lines representative of the endometriotic lesion microenvironment (**Figure 4**). Specifically, selected cell lines were representative of endometriotic epithelial (12Z) and endometrial stromal (hESC) cells, as well as vasculature/endothelial cells (HUVEC) and endometrial epithelial cells (EECC). Multiplex cytokine array of supernatant produced by rhIL-23 treated 12Z, EECC, HUVEC, and hESC cell lines revealed that IL-23 treatment significantly altered the production of platelet-derived growth factor (PDGF)-AA and PDGF-AB/BB by 12Z, EECC, and HUVEC cell lines (**Fig. 4A-C**), as well as IFNγ-inducible protein 10 (IP-10) and fibroblast growth factor-2 (FGF-2) production by EECC and HUVEC cell lines, respectively (**Fig. 4E-F**). While hESC cells also produced notable PDGF levels in response to IL-23 treatment, there were no significant differences found between treatment groups (data not shown). Additionally, rhIL-23 treatment had dose-dependent effects on hESCs, with significant increases in both IL-1β (**Fig. 4D**) and Fms-related tyrosine kinase 3 ligand (FLT-3L; **Fig. 4G**) as IL-23 dosage increased.

**Figure 4:**
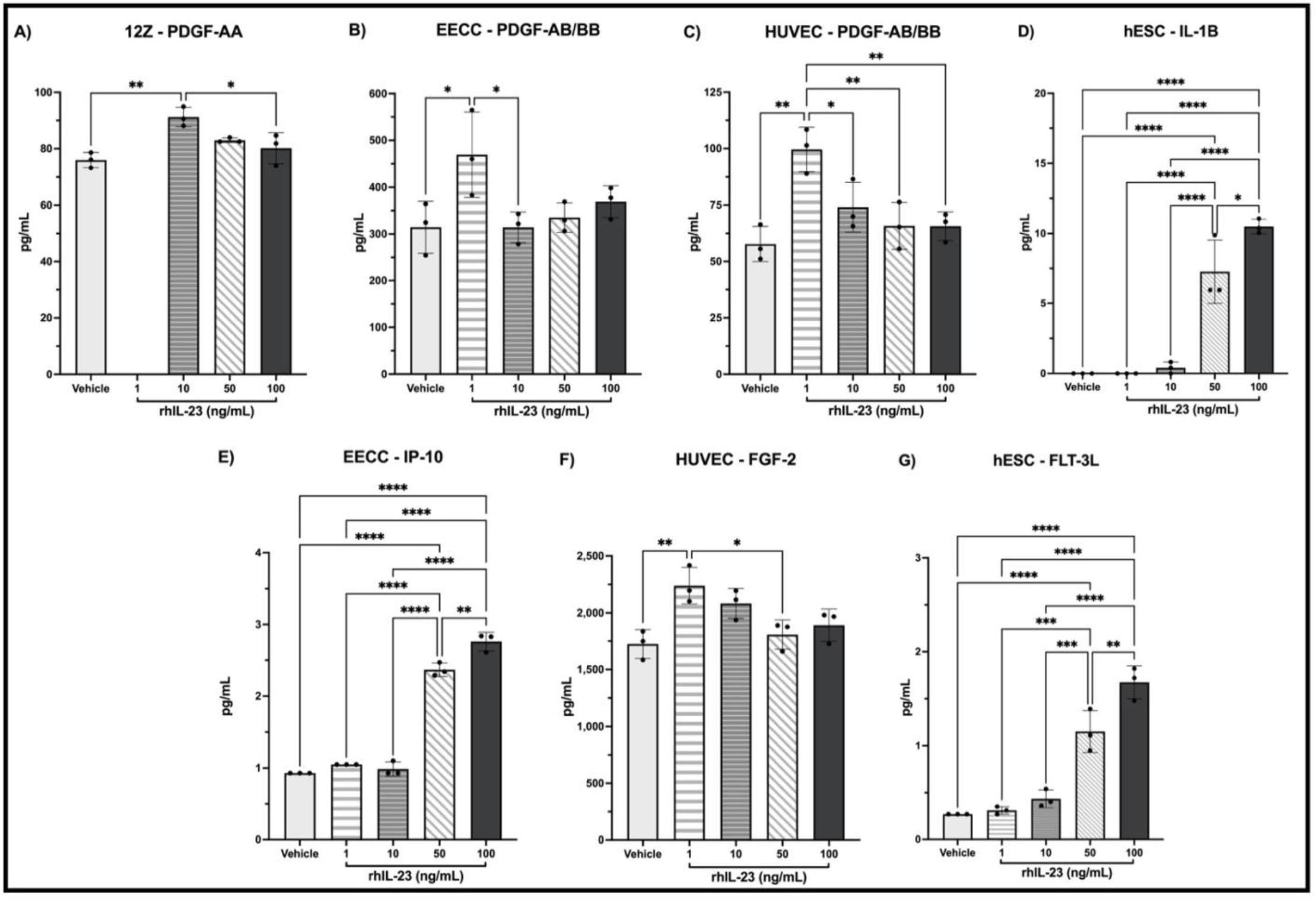
rhIL-23 stimulation of cells representative of the endometriotic microenvironment significantly alter the production of cytokines/chemokines known to play a role in lesion establishment and maintenance. 12Z, EECC, HUVEC, and hESC cell lines were stimulated in triplicates for 24hrs with 0.01% BSA (bovine serum albumin) in PBS (vehicle) or varying concentrations of rhIL-23 (1, 10, 50, 100ng/mL) prior to supernatant collection and analysis via multiplex cytokine array. 48 analytes were measured with focus on inflammatory and angiogenic factors, as well as mediators in the T_H_17 pathway. Most notably, IL-23 treatment significantly altered the production of PDGF by 12Z, EECC, and HUVEC cells (**A-C**). In a dose-dependent fashion, IL-23 stimulation significantly increased IL-1β and FLT-3L production by hESCs (**D, G**), IP-10 production by EECCs (**E**). IL-23 treatment also significantly increased FGF-2 levels in HUVECs (**F**). Data is represented as mean ± SD. A one-way ANOVA with Tukey post-hoc was used to assess significance. *p<0.05, **p<0.01, ***p<0.001, ****p<0.0001.

To assess the influence of IL-23 on cell proliferation and apoptosis we conducted WST-1 and caspase 3/7 glo assays respectively (**Supplementary Fig. 2**). We found that rhIL-23 treatment significantly increased hESC cell proliferation in the 1ng/mL and 100ng/mL treatment groups, though there were no significant differences in the proliferation of 12Z, EECC, or HUVEC cells.

There were also no significant differences in cell death of 12Zs, EECCs, HUVECs, or hESCs following IL-23 treatment. Moreover, we sought to determine the influence of IL-23 on angiogenesis using a tubulogenesis assay with HUVECs. Though, no significant differences were found in tube length or number of total branching points when comparing rhIL-23 treated groups to the PBS control (**Supplementary Fig. 3**).

### IL-23 containing T_H_17 derivation cocktail significantly drives pathogenic T_H_17 cell fate in mice

Literature shows that IL-23 signalling is necessary to drive, maintain, and expand a pathogenic T_H_17 cell fate^18,20,43^. Thus, we sought to model this *in-vitro*, examining whether IL-23 alters production of pathogenic T_H_17 cells from naïve CD4^+^ T cells isolated from murine splenocytes via flow cytometry (**Figure 5A**). Results show that treatment with an IL-23 containing T_H_17 derivation cocktail significantly increased frequency and number of pathogenic T_H_17 cells from murine naïve CD4^+^ T cells as compared to all other treatment groups, as IL-23 with activation and media with activation groups were not able to drive pathogenic T_H_17 cells (**Fig. 5B-C**).

**Figure 5:**
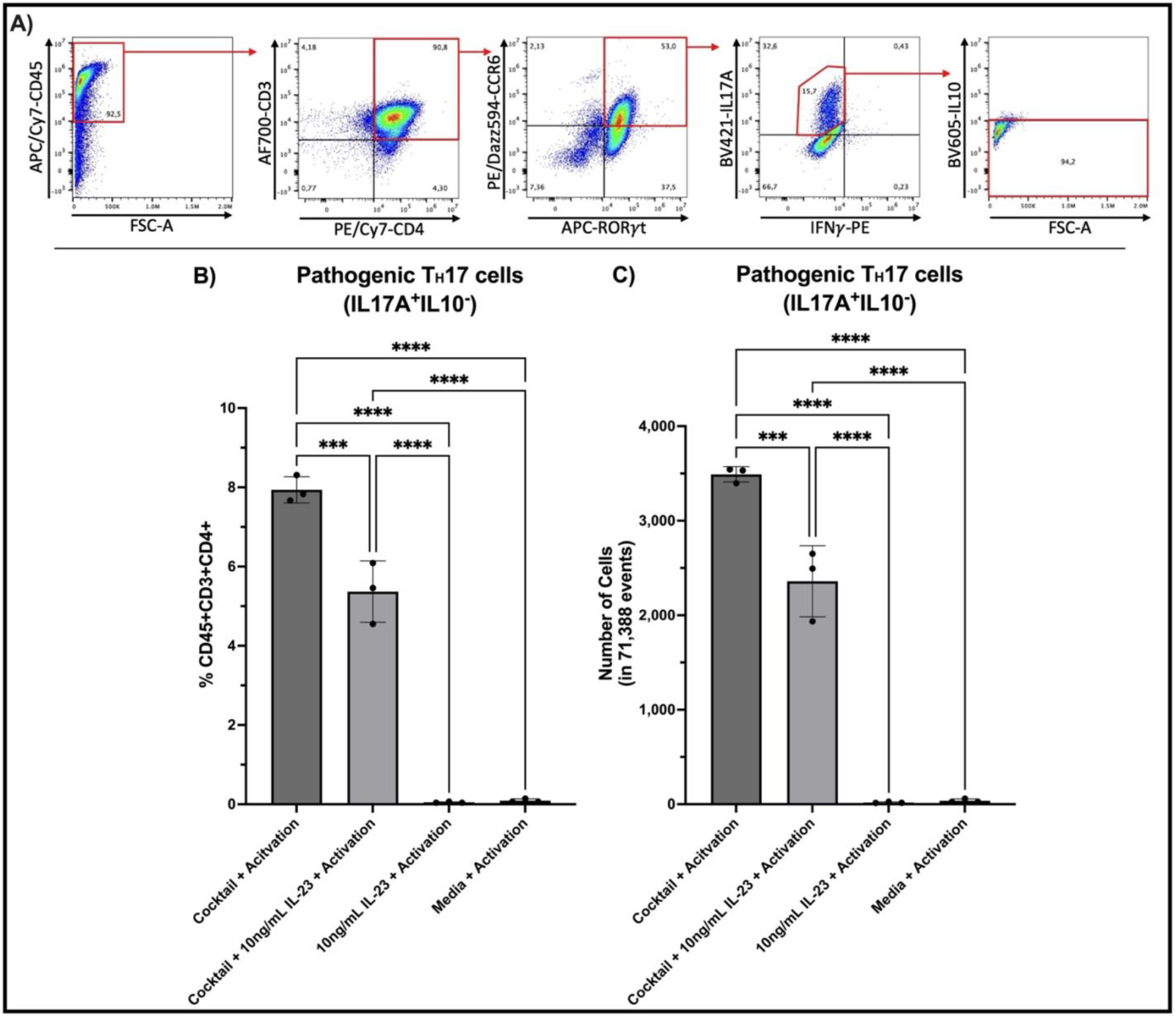
T_H_17 derivation cocktail significantly promotes the production of pathogenic T_H_17 cells from murine naïve CD4^+^ T cells. Naïve CD4^+^ T cells isolated from murine splenocytes were treated in-vitro in triplicates for 5 days with the cocktail and CD3/CD28 activation, cocktail and 10ng/mL rmIL-23 with activation, 10ng/mL rmIL-23 and activation, or media with activation. Cells were re-stimulated with PMA/ionomycin and protein transport inhibitor cocktail prior to flow cytometric analysis for pathogenic T_H_17 cells (CD45^+^CD3^+^CD4^+^CCR6^+^RORγt^+^IL17A^+^IFNγ^-^IL10^−^; **A**). Naïve CD4^+^ T cells treated with the T_H_17 derivation cocktail and activation produced a significantly higher frequency of pathogenic T_H_17 cells within the CD45^+^CD3^+^CD4^+^ population (**B**) as well as number of T_H_17 cells per 71,388 events (**C**), as compared to all other treatment groups. Number of T_H_17 cells was normalized to the lowest acquired events amongst all samples (71,388 events) using FlowJo software. Data is represented as mean ± SD. A one-way ANOVA with Tukey post-hoc was used to assess significance. ***p<0.001, ****p<0.0001.

**Figure 6:**
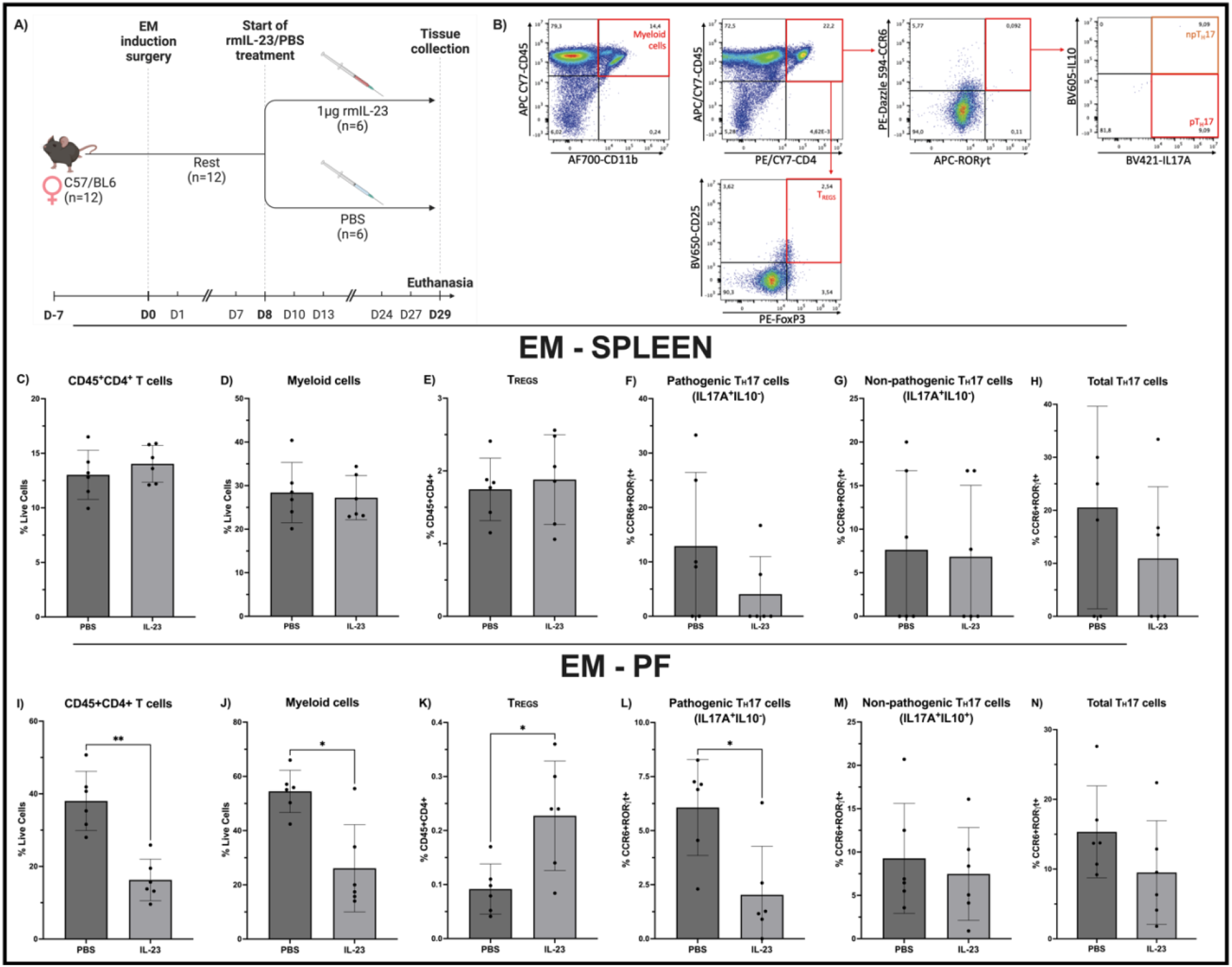
IL-23 treatment results in significant immune dysfunction localized to the murine endometriotic microenvironment, confirmed via flow cytometry. 1 week following surgery, endometriosis (EM)-induced mice were treated (i.p.) with 1μg rmIL-23 (n=6) or PBS (n=6) 3 times a week for 3 weeks before splenocytes and PF were isolated for flow cytometric analysis (**A**). Markers were used to detect CD4^+^ T cells (CD45^+^CD4^+^), myeloid cells (CD45^+^CD11b^+^), T_REGS_ (CD45^+^CD4^+^CD25^+^FoxP3^+^), pathogenic T_H_17 cells (pT_H_17; CD45^+^CD4^+^CCR6^+^RORγt^+^IL17A^+^IL10^−^), non-pathogenic T_H_17 cells (npT_H_17; CD45^+^CD4^+^CCR6^+^RORγt^+^IL17A^+^IL10^+^), and total T_H_17 cells (CD45^+^CD4^+^CCR6^+^RORγt^+^IL17A^+^; **B**). Results depict respective immune cell populations within the spleen (**C-H**) as well as the PF (**I-N**). Data is represented as mean ± SD. A non-parametric student’s t-test with Mann-Whitney correction was used to assess significance. *p<0.05, **p<0.01. Graphic adapted from “Mouse High Fat Diet Experimental Timeline”, by BioRender.com (2023). Retrieved from https://app.biorender.com/biorender-templates.

### IL-23 treatment significantly dysregulates the local immune microenvironment in a mouse model of endometriosis

To assess the influence of IL-23 on immune cell recruitment/immune dysfunction in endometriosis, C57BL/6 mice (n=12) underwent endometriosis induction surgery and mice were treated with rmIL-23 (n=6) or PBS (controls, n=6) via i.p. injections three times a week for three weeks. Murine splenocytes and PF were collected to assess systemic and local immune alterations, respectively, via flow cytometry (**Figure 6**). Results depicted significant alterations to CD4^+^ T cells, myeloid cells, T_REGS_, and pathogenic T_H_17 cells in the PF, though there were no significant differences in immune cell subsets within murine splenocytes. As speculated, IL-23 significantly altered the T_H_17/T_REG_ axis in murine PF, increasing T_REGS_ and decreasing pathogenic T_H_17 cells (**Fig. 6J-K**), though these differences were not statistically significant in murine splenocytes (**Fig. 6D-E**). The significant differences seen in immune cell subsets within the PF were not detected in our sham mouse model (**Supplementary Fig. 4**), highlighting the influence of IL-23 specifically in the context of endometriosis. IL-23 was also found to significantly increase immune cell recruitment to the peritoneal cavity, irrespective of disease context (endometriosis/sham) (**Supplementary Fig. 5**).

In addition to inflammation, lesion proliferation and vascularization are known to be hallmark features of endometriosis pathophysiology^6,44,45^. Thus, we aimed to examine the role of IL-23 on lesion vascularization and proliferation using an established mouse model of endometriosis (**Figure 7**). H&E staining confirmed that murine endometriotic lesions were representative of endometriosis lesion histology (**Fig. 7A-B**). Additionally, lesions from IL-23 treated mice had large, multinucleated cells consistent with the histology of giant cells (**Fig. 7C**). To our knowledge, this is the first report identifying giant cells within murine endometriosis-like lesions. Though these cells were also found in some PBS treated mice, giant cells were significantly increased in the IL-23 treated group (**Fig. 7D**). Lesions were also subjected to IHC for markers of proliferation (Ki67) and vascularization (CD31; **Fig. 7E-F, H-I**). IL-23 treated mice had trends of increased lesion proliferation and vascularization, though differences were not statistically significant (**Fig. 7G, J**).

**Figure 7:**
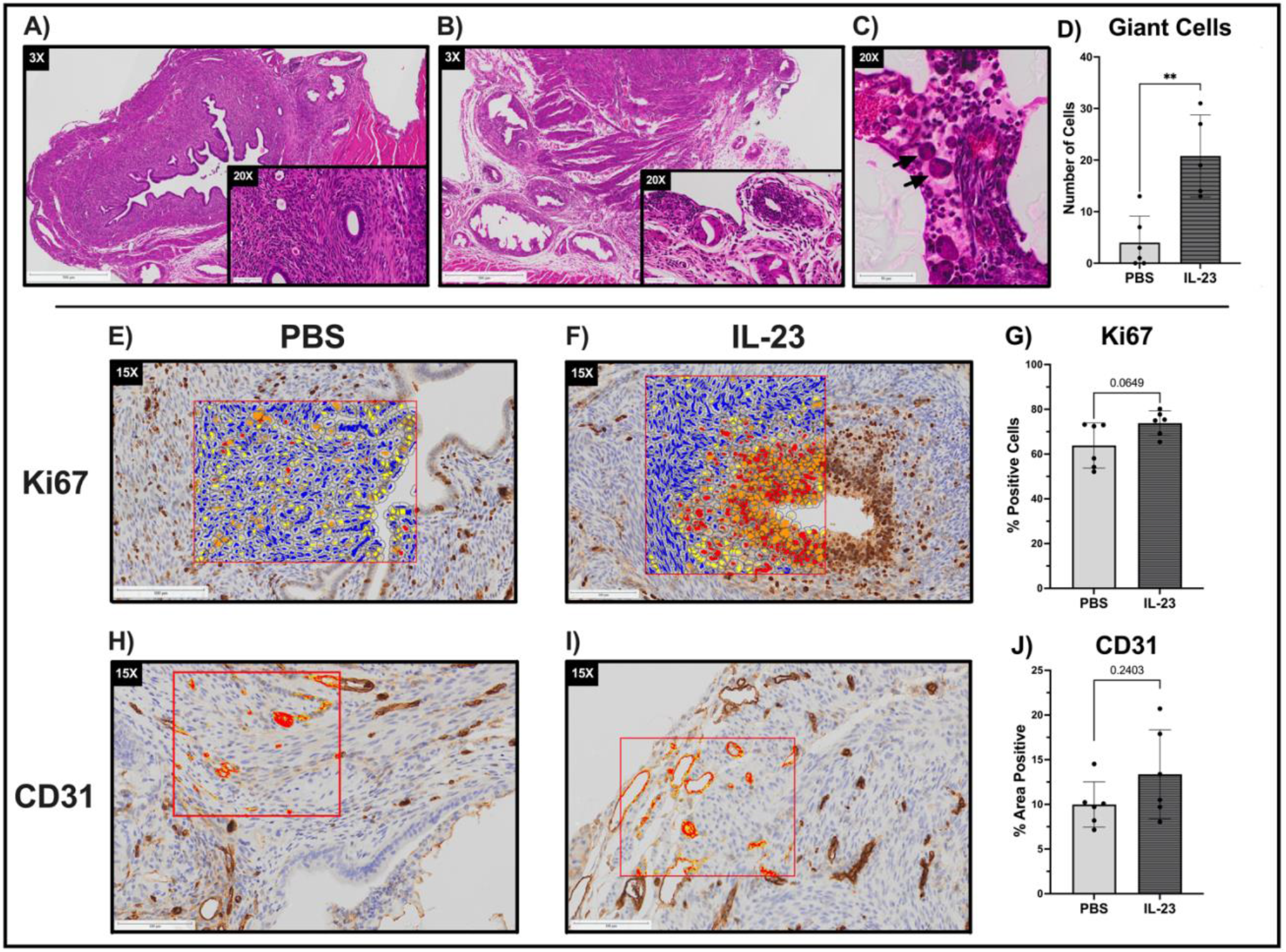
IL-23 treatment results in significantly increased numbers of giant cells within lesions as well as trends of increased lesion proliferation and vascularization. 1 week following endometriosis induction surgery, mice were administered (i.p.) 1μg rmIL-23 (n=6) or PBS (n=6) 3 times a week for 3 weeks. The lesions of both PBS (**A**) and rmIL-23 (**B**) treated mice well recapitulate endometriosis lesion histology, as shown by H&E staining. rmIL-23 treated mice had significantly increased numbers of large, multinucleated cells within their lesions consistent with giant cell histology, as indicated by black arrows (**C-D**). Murine lesions were subjected to IHC for markers of proliferation (Ki67; **E-F**) and vascularization (CD31; **H-I**). A cytonuclear algorithm was used to detect percentage of cells expressing Ki67 as compared to the total cell population (**G**), while percent area quantification was used to analyze vascularization (**J**). rmIL-23 treated mice had trends of increased proliferation and vascularization, though these differences were not statistically significant. Statistical significance was assessed using a non-parametric student’s t-test with Mann-Whitney correction. **p<0.01. Scanned H&E images were digitally analyzed using HALO Imaging Software (Indica Labs, USA) and provided at 3x and 20x magnification; scale bars 500μm and 50μm respectively. Similarly, scanned IHC images are provided at 15x magnification; scale bar 100μm.

**Figure 8:**
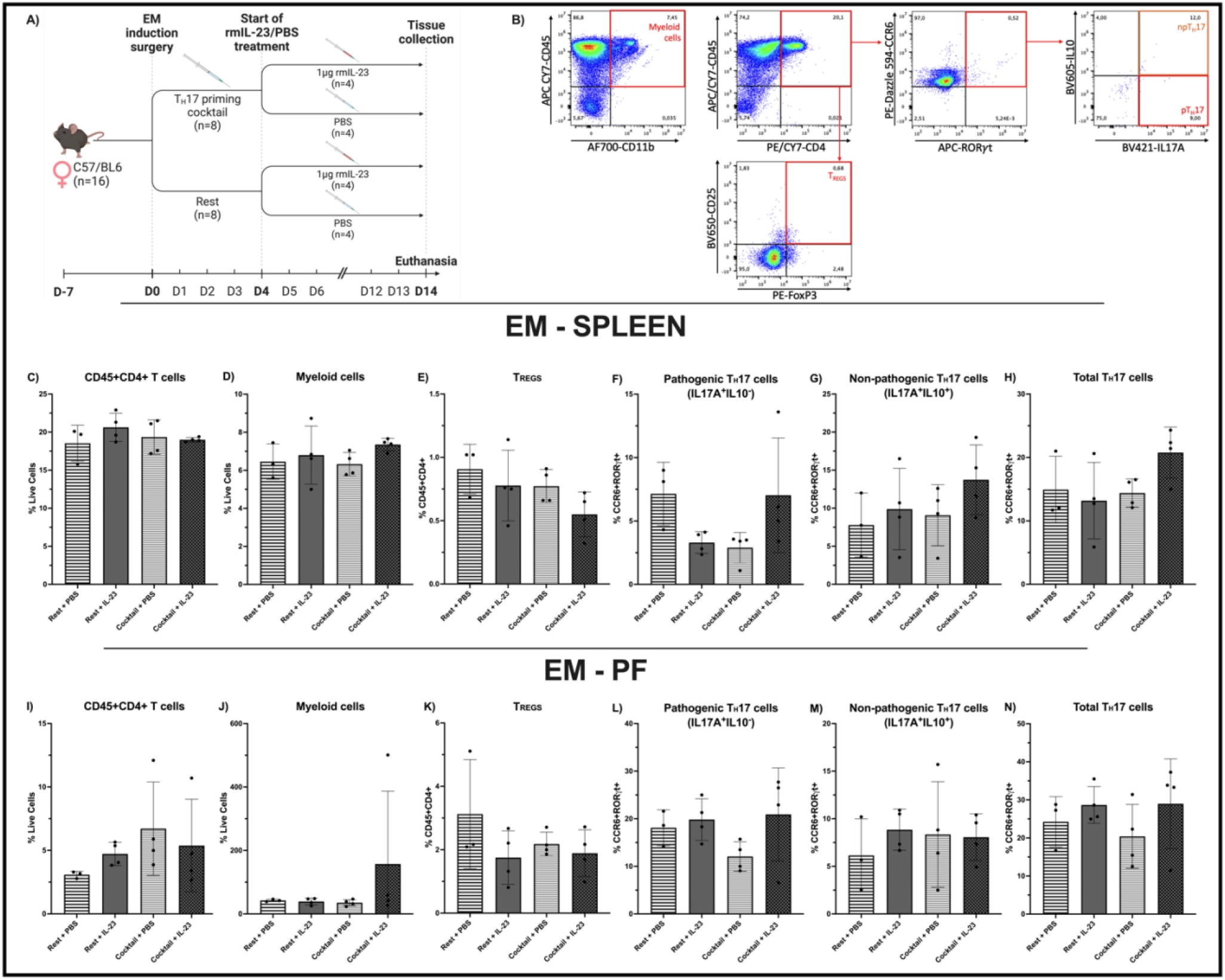
IL-23 and/or T_H_17 priming cocktail do not significantly alter local or systemic immune response during a short time course study in a mouse model of endometriosis. Following endometriosis (EM) induction surgery in C57BL/6 mice (n=16), mice were rested for 4 days (n=8) or treated daily (i.p.) with a T_H_17 priming cocktail (0.5μg rmIL-6, 1μg rmTGF-β1; n=8). Following this, half of the mice in each cohort were treated daily (i.p.) for a further 10 days with either 1μg rmIL-23 (n=4) or PBS (controls; n=4) prior to euthanasia (**A**). Markers were used to detect CD4^+^ T cells (CD45^+^CD4^+^), myeloid cells (CD45^+^CD11b^+^), T_REGS_ (CD45^+^CD4^+^CD25^+^FoxP3^+^), pathogenic T_H_17 cells (pT_H_17; CD45^+^CD4^+^CCR6^+^RORγt^+^IL17A^+^IL10^−^), non-pathogenic T_H_17 cells (npT_H_17; CD45^+^CD4^+^CCR6^+^RORγt^+^IL17A^+^IL10^+^), and total T_H_17 cells (CD45^+^CD4^+^CCR6^+^RORγt^+^IL17A^+^; **B**). Results depict respective immune cell populations within the spleen (**C-H**) as well as the PF (**I-N**). Data is represented as mean ± SD. No statistically significant differences were found, as assessed by a one-way ANOVA with Tukey post-hoc. Graphic adapted from “Mouse High Fat Diet Experimental Timeline”, by BioRender.com (2023). Retrieved from https://app.biorender.com/biorender-templates.

IL-23 is known to promote inflammation and pathogenic T_H_17 cell fate in combination with other inflammatory cytokines^18,46^. Thus, we sought to determine if treatment with a T_H_17 priming cocktail and/or rmIL-23 influenced the production of pathogenic T_H_17 cells, and in-turn the inflammatory milieu, within a mouse model of endometriosis (**Figure 8**). Splenocytes and PF were collected and analyzed via flow cytometry, however, no statistically significant differences in immune cell subsets were detected in treated groups as compared to controls.

## Discussion

IL-17, a proinflammatory and proangiogenic cytokine known to be largely secreted by T_H_17 cells, has been linked in the pathogenesis of endometriosis^10^. While exact mechanisms are unknown, production of IL-17, as well as other downstream cytokines by T_H_17 cells, may exacerbate endometriosis development by recruiting immune cells to the lesion site, such as neutrophils, macrophages, and lymphocytes, which collectively function to enhance lesion establishment^12,28^.

Specifically, IL-17 is increased in patient plasma and PF and this is correlated with disease severity^47^. Additionally, IL-17 is produced by lesions and following excision surgery, a significant reduction of systemic IL-17 levels was reported^10^, highlighting the potential link between IL-17 and endometriosis. Furthermore, T_H_17 cells are dysregulated in endometriosis^24,29,33^ and are also associated with disease severity^13^. Thus, while this downstream cytokine in the T_H_17 axis has been examined in endometriosis, there is a gap in knowledge as to what is driving T_H_17/IL-17 axis dysregulation in endometriosis. IL-23 plays a key role in governing T_H_17 cell fate^20,48^, driving pathogenic T_H_17 cells producing IL-17 and downregulating anti-inflammatory IL-10^17,18^. Ultimately, as emerging evidence has underscored the role of IL-23 in driving this axis to produce IL-17 in various other diseases^30,49^, we sought to examine if IL-23 may also be driving the T_H_17/IL-17 axis to exacerbate disease in endometriosis.

Our studies using patient samples reveal significant dysregulation in the IL-23/T_H_17 axis in patient tissues as compared to healthy controls. Specifically, RT qPCR results show *IL17A, IL17F, IL21, IL22, IL12B* (IL-23 p40 subunit), and *TGFB1* genes to be significantly upregulated in ectopic tissues as compared to matched eutopic tissue and control samples. This is substantial as IL-17A, IL-17F, IL-21, and IL-22 are downstream cytokines produced by T_H_17 cells, while IL-23 and TGF-β1 are important in driving and maintaining a pathogenic T_H_17 cell fate^20,48^. RT qPCR results also suggest patients may have increased sensitivity to IL-23 signalling, as *IL23R* was significantly upregulated in both ectopic and eutopic patient tissues as compared to controls. Moreover, *IL12B*, encoding the p40 subunit shared between IL-23 and IL-12, was significantly increased in accordance with literature^50^. This increase is unlikely to indicate IL-12 involvement as the p35 subunit (*IL12A*), specific to IL-12, was not significantly increased. Instead, this may indicate involvement of p40 homodimers or p40 monomers, which are increased in endometriosis PF^51^.

Complimentary to the RT qPCR results, IL-23 protein was detected in endometriotic lesions and was significantly increased in patient plasma compared to healthy controls. Though IL-23 protein expression was not significantly different across patient ectopic (ovarian endometrioma) or eutopic tissues as compared to controls, IL-23 was significantly increased in luminal and glandular epithelium as compared to stroma. As endometrioma lesions are more cystic in nature, they are comprised of less luminal and glandular epithelium compared to non-cystic lesions. Thus, as we illustrate IL-23 to be predominately localized to epithelial tissue, IL-23 protein expression is likely reduced in endometrioma lesions compared to other lesion subtypes. Though, further research is required to quantify differential IL-23 protein expression across lesion subtypes. Others do report significantly increased IL-23 protein in endometriosis patient PF, ectopic lesions, and serum, which is correlated with increased levels of IL-17^52–55^. Moreover, IL-23 is significantly increased in both patient follicular fluid and serum in later stages of disease (III-IV) compared to earlier stages (I-II)^53^. Highlighting that IL-23 may be associated with disease severity. Contrary to this, some studies report no significant differences in IL-23 in patient serum when not stratified by disease stage^53,54^. While we did not capture significant differences in *IL23A* (p19 subunit) in patient tissues at the mRNA level, dysregulation of other key members within IL-23/T_H_17 axis combined with significantly increased IL-23 protein levels in patient plasma indicate involvement in endometriosis pathophysiology.

In our *in-vitro* studies, IL-23 cocktail treatment in combination with CD3/CD28 activation significantly promoted T_H_17 cell differentiation from human and murine naïve CD4^+^ T cells. This cocktail was selected as evidence shows the endometriosis milieu contains and is influenced by TGF-β1, IL-1β, IL-6, IL-21, and IL-23 cytokines^53,56–59^, which are also vital in driving a pathogenic T_H_17 cell fate^17,60^. Thus, we aimed to partially recapitulate this environment *in-vitro*. Results depict that naïve human CD4^+^ T cells driven in the presence of this combination treatment produce significantly increased IL-17A/F and decreased IL-10, characteristic of pathogenic T_H_17 cells. This confirms that T_H_17 cells exposed to IL-23 cocktail acquire a pathogenic phenotype consistent with literature^12,20,48^. Additionally, human naïve CD4^+^ T cells driven with this combination treatment produce significantly increased MCP-1, CCL5, CXCL9, and MDC. As these factors play important chemotactic roles, IL-23-driven and activated T_H_17 cells may promote immune cell recruitment and produce proinflammatory factors, such as IL-17, to further exacerbate lesion inflammation, proliferation, and angiogenesis. However, while there are reports of pathogenic T_H_17 cells producing CCL5^12^, it is unclear to what abundance T_H_17 cells may produce MCP-1, CXCL9, and MDC. Further studies are required to examine co-production of these factors by T_H_17 cells.

As in other pathologies IL-23 is produced in response to damage-associated molecular patterns (DAMPs), and literature suggests DAMPs/alarmins are released during endometriosis lesion establishment^57^, IL-23 may not only act in endometriosis by promoting and supporting pathogenic T_H_17 cell fate, but also may directly act on the lesion to perpetuate inflammatory signals in response to DAMPs. Thus, we sought to determine the independent effects of IL-23 treatment on cells representative of the endometriotic microenvironment, specifically 12Z, EECC, HUVEC, and hESC cells. Results showed that *in-vitro* rhIL-23 stimulation of cell lines significantly altered the production of PDGF-AA from 12Zs, PDGF-AB/BB levels from both EECCs and HUVECs, as well as FGF-2 production from HUVECs. Additionally, *in-vitro* rhIL-23 treatment produced significant dose-dependent increases of IP-10 by EECCs and IL-1β and FLT-3L by hESCs. rhIL-23 treatment also significantly increased proliferation of hESCs, a major component of the endometriotic lesion. Collectively, our *in-vitro* data suggests that IL-23 stimulation significantly alters the production of factors known to play a role in lesion establishment and maintenance and may promote lesion growth/survival.

Our mouse model of endometriosis has been extensively used to gain mechanistic insights into the complex pathophysiology of endometriosis in previously reported studies^34,61–63^. Indeed, we identify significantly increased numbers of large, multinucleated cells in lesions obtained from IL-23 treated mice as compared to controls (PBS). Histologically, these cells appear to be giant cells of monocyte origin, however further characterization is necessary. While we captured trends of increased lesion proliferation and vascularization in IL-23 treated mice as compared to controls, these differences were not statistically significant. This may be reflective of our mouse model, the time course, and/or the dose, and route of IL-23 administration, as IL-23 is known to have effects on cell proliferation and vascularization^64–67^. We also used this mouse model to determine if IL-23 treatment alters systemic and/or local immune cell subsets, and in-turn, may promote immune dysfunction associated with endometriosis. Indeed, IL-23 treatment significantly reduced CD4^+^ T cells and myeloid cells in murine PF as compared to controls, possibly due to homing of CD4^+^ T cells and myeloid cells to the lesion. Additionally, IL-23 treatment dysregulated the local T_H_17/T_REG_ axis, where pathogenic T_H_17 cells were significantly reduced and T_REGS_ were significantly increased in the PF of mice treated with IL-23. This T_H_17/T_REG_ axis dysregulation is noteworthy as balance between T_H_17 and T_REGS_ is necessary to maintain immune homeostasis and avoid dysfunction^23,68–70^. These significant alterations in immune cell subsets within murine PF were not observed in our sham mouse model, highlighting that these differences were specific to an endometriosis-like microenvironment. Finally, as literature suggests TGF-β1 and IL-6 are required for initial derivation of T_H_17 cells, while IL-23 is necessary for full induction and maintenance of pathogenic T_H_17 cell fate^71,72^, we used a mouse model to assess the influence of IL-23 alone and in combination with a T_H_17 priming cocktail (TGF-β1 and IL-6) during endometriosis lesion establishment. Our short time course study revealed no significant differences in murine immune cell subsets between treatment groups. Thus, further research is required to establish the specific combination of cytokines and timeframe necessary to capture murine T_H_17 differentiation *in-vivo*. Our *in-vivo* data reveals that IL-23 treatment significantly influences local immune dysfunction in endometriosis and promotes immune cell recruitment to the peritoneal cavity, which may exacerbate local immune dysfunction to promote lesion survival and the well-known inflammatory milieu in endometriosis.

In conclusion, while our data suggests that the IL-23/T_H_17 axis is dysregulated in endometriosis, it is important to note that there is no existing evidence in literature mechanistically linking this axis to endometriosis pathology. We acknowledge there are limitations to this study and those investigating endometriosis in general due to limited access in obtaining sufficient numbers of well categorized patient samples stratified by disease stage, subtype, and menstrual cycle, as patients commonly experience irregular menstruation. The diagnostic challenges of endometriosis also complicate this due to invasive diagnostic measures, heterogenous symptom presentation, normalization of patient symptoms, and stigmatization of reproductive disorders. This commonly results in long diagnostic delays and misdiagnosis of patients. Due to limitations in obtaining fresh patient samples, we were unable to isolate primary cells from endometriosis lesions, and thus we acknowledge the known limitations of using immortalized cell lines. However, we do supplement this data from immortalized cell lines with that from primary human CD4^+^ T cells. While we use our syngeneic immunocompetent mouse model of endometriosis to provide insights into the role of IL-23 in endometriosis pathophysiology, this model is not able to completely recapitulate human disease as rodents do not spontaneously develop endometriosis. Having said this, we report significant dysregulation of the IL-23/T_H_17 axis in endometriosis patients, and this was captured in *in-vitro* studies and our *in-vivo* mouse model. As there is currently no effective long-term, non-invasive, or non-hormonal therapy option existing for endometriosis patients who aim to conceive, there is an urgent need to explore novel therapeutics for endometriosis. Ultimately, IL-23 may provide a promising therapeutic avenue to reduce the large burden of endometriosis, as IL-23 targeting therapeutics have been approved for human use and/or are currently in development for various chronic inflammatory diseases, including rheumatoid arthritis and psoriasis^32,73–77^.

## Author Contributions

DS conceived and conducted experiments, analyzed data, and wrote the manuscript. KZ, JM, HL, SH, and AM conducted experiments. OB and BL contributed human patient samples. CT contributed reagents, conceived experiments, provided financial support. All authors read and edited the manuscript.

## Acknowledgements

We would like to thank Oliver Jones at Queen’s CardioPulmonary Unit (QCPU, Queen’s University) for all histology processing. We also thank Wei Wang at Kingston Health Sciences Centre (KHSC, Queen’s University) for all immunohistochemistry processing. Finally, we thank Xiao Zhang at Queen’s Laboratory for Molecular Pathology (QMLP, Queen’s University) for aid with slide scanning.

## References

1. Giudice, L. C. Endometriosis. N. Engl. J. Med. 362, 2389–2398 (2010).

2. Alimi, Y., Iwanaga, J., Loukas, M. & Tubbs, R. S. The Clinical Anatomy of Endometriosis: A Review. Cureus (2018). doi:10.7759/cureus.3361

3. Allaire, C., Bedaiwy, M. A. & Yong, P. J. Diagnosis and management of endometriosis. Can. Med. Assoc. J. 195, E363–E371 (2023).

4. Shafrir, A. L. et al. Risk for and consequences of endometriosis: A critical epidemiologic review. Best Pract. Res. Clin. Obstet. Gynaecol. 51, 1–15 (2018).

5. Horne, A. W. & Missmer, S. A. Pathophysiology, diagnosis, and management of endometriosis. BMJ e070750 (2022). doi:10.1136/bmj-2022-070750

6. Zondervan, K. T. et al. Endometriosis. Nat. Rev. Dis. Prim. 4, 9 (2018).

7. Osuga, Y. et al. Th2 cells and Th17 cells in the development of endometriosis – possible roles of interleukin-4 and interleukin-17A. Journal of Endometriosis 8, (2016).

8. Miossec, P. & Kolls, J. K. Targeting IL-17 and TH17 cells in chronic inflammation. Nat. Rev. Drug Discov. 11, 763–776 (2012).

9. Bartlett, H. S. & Million, R. P. Targeting the IL-17–TH17 pathway. Nat. Rev. Drug Discov. 14, 11–12 (2015).

10. Ahn, S. H. et al. IL-17A Contributes to the Pathogenesis of Endometriosis by Triggering Proinflammatory Cytokines and Angiogenic Growth Factors. J. Immunol. 195, (2015).

11. Patel, D. D. & Kuchroo, V. K. Th17 Cell Pathway in Human Immunity: Lessons from Genetics and Therapeutic Interventions. Immunity 43, 1040–1051 (2015).

12. Wu, X., Tian, J. & Wang, S. Insight Into Non-Pathogenic Th17 Cells in Autoimmune Diseases. Front. Immunol. 9, (2018).

13. Gogacz, M. et al. Increased percentage of Th17 cells in peritoneal fluid is associated with severity of endometriosis. J. Reprod. Immunol. 117, (2016).

14. Stockinger, B. & Omenetti, S. The dichotomous nature of T helper 17 cells. Nat. Rev. Immunol. 17, (2017).

15. Hirota, K. et al. Plasticity of TH17 cells in Peyer’s patches is responsible for the induction of T cell–dependent IgA responses. Nat. Immunol. 14, (2013).

16. Conti, H. R. et al. Oral-resident natural Th17 cells and γδ T cells control opportunistic Candida albicans infections. J. Exp. Med. 211, (2014).

17. McGeachy, M. J. et al. TGF-β and IL-6 drive the production of IL-17 and IL-10 by T cells and restrain TH-17 cell–mediated pathology. Nat. Immunol. 8, 1390–1397 (2007).

18. Lee, Y. et al. Induction and molecular signature of pathogenic TH17 cells. Nat. Immunol. 13, 991–999 (2012).

19. Teng, M. W. L. et al. IL-12 and IL-23 cytokines: from discovery to targeted therapies for immune-mediated inflammatory diseases. Nat. Med. 21, 719–729 (2015).

20. Langrish, C. L. et al. IL-23 drives a pathogenic T cell population that induces autoimmune inflammation. J. Exp. Med. 201, 233–240 (2005).

21. Ma, R., Su, H., Jiao, K. & Liu, J. Role of Th17 cells, Treg cells, and Th17/Treg imbalance in immune homeostasis disorders in patients with chronic obstructive pulmonary disease. Immunity, Inflamm. Dis. 11, (2023).

22. Wang, R. et al. Neutralizing IL-23 Is Superior to Blocking IL-17 in Suppressing Intestinal Inflammation in a Spontaneous Murine Colitis Model. Inflamm. Bowel Dis. 21, 973–984 (2015).

23. Lee, G. The Balance of Th17 versus Treg Cells in Autoimmunity. Int. J. Mol. Sci. 19, 730 (2018).

24. Khan, K. N. et al. Differential Levels of Regulatory T Cells and T-Helper-17 Cells in Women With Early and Advanced Endometriosis. J. Clin. Endocrinol. Metab. 104, 4715–4729 (2019).

25. Maloy, K. J. & Kullberg, M. C. IL-23 and Th17 cytokines in intestinal homeostasis. Mucosal Immunol. 1, 339–349 (2008).

26. Vallvé-Juanico, J., Houshdaran, S. & Giudice, L. C. The endometrial immune environment of women with endometriosis. Hum. Reprod. Update 25, 565–592 (2019).

27. Hirata, T. et al. Interleukin (IL)-17A Stimulates IL-8 Secretion, Cyclooxygensase-2 Expression, and Cell Proliferation of Endometriotic Stromal Cells. Endocrinology 149, 1260–1267 (2008).

28. Miller, J. E. et al. IL-17A Modulates Peritoneal Macrophage Recruitment and M2 Polarization in Endometriosis. Front. Immunol. 11, (2020).

29. Miller, J. E. et al. T helper 17 axis and endometrial macrophage disruption in menstrual effluent provides potential insights into the pathogenesis of endometriosis. F&S Sci. (2022). doi:10.1016/j.xfss.2022.04.007

30. Schinocca, C. et al. Role of the IL-23/IL-17 Pathway in Rheumatic Diseases: An Overview. Front. Immunol. 12, (2021).

31. Iwakura, Y. & Ishigame, H. The IL-23/IL-17 axis in inflammation. J. Clin. Invest. 116, (2006).

32. Torres, T. Selective Il-23 Inhibitors: The New Kids on the Block in the Treatment of Psoriasis. Actas Dermo-Sifiliográficas (English Ed. 109, (2018).

33. Takamura, M. et al. Simultaneous Detection and Evaluation of Four Subsets of CD4+ T Lymphocyte in Lesions and Peripheral Blood in Endometriosis. Am. J. Reprod. Immunol. 74, 480–486 (2015).

34. Zutautas, K. B. et al. The dysregulation of leukemia inhibitory factor and its implications for endometriosis pathophysiology. Front. Immunol. 14, (2023).

35. Symons, L. K. et al. Neutrophil recruitment and function in endometriosis patients and a syngeneic murine model. FASEB J. 34, 1558–1575 (2020).

36. Kleinewietfeld, M. et al. Sodium chloride drives autoimmune disease by the induction of pathogenic TH17 cells. Nature 496, 518–522 (2013).

37. Delens, L. et al. In Vitro Th17-Polarized Human CD4+ T Cells Exacerbate Xenogeneic Graft-versus-Host Disease. Biol. Blood Marrow Transplant. 25, 204–215 (2019).

38. Esplugues, E. et al. Control of TH17 cells occurs in the small intestine. Nature 475, 514–518 (2011).

39. Bettelli, E. et al. Reciprocal developmental pathways for the generation of pathogenic effector TH17 and regulatory T cells. Nature 441, 235–238 (2006).

40. Wimasis. (2016).

41. Cotechini, T., Jones, O. & Hindmarch, C. C. T. Imaging Mass Cytometry in Immuno-Oncology. in 1–15 (2023). doi:10.1007/978-1-0716-2914-7_1

42. Veldhoen, M., Hocking, R. J., Atkins, C. J., Locksley, R. M. & Stockinger, B. TGFβ in the Context of an Inflammatory Cytokine Milieu Supports De Novo Differentiation of IL-17-Producing T Cells. Immunity 24, 179–189 (2006).

43. Meyer zu Horste, G. et al. RBPJ Controls Development of Pathogenic Th17 Cells by Regulating IL-23 Receptor Expression. Cell Rep. 16, 392–404 (2016).

44. Laschke, M. W. & Menger, M. D. Basic mechanisms of vascularization in endometriosis and their clinical implications. Hum. Reprod. Update 24, 207–224 (2018).

45. Tayade, C. S11.2: Role of endocannabinoids in endometriosis. Am. J. Reprod. Immunol. 87, 55–55 (2022).

46. Manel, N., Unutmaz, D. & Littman, D. R. The differentiation of human TH-17 cells requires transforming growth factor-β and induction of the nuclear receptor RORγt. Nat. Immunol. 9, 641–649 (2008).

47. Zhang, X., Xu, H., Lin, J., Qian, Y. & Deng, L. Peritoneal fluid concentrations of interleukin-17 correlate with the severity of endometriosis and infertility of this disorder. BJOG An Int. J. Obstet. Gynaecol. 112, 1153–1155 (2005).

48. Kleiner, J. C. & Krebs, C. F. Persistence, Pathogenicity and Plasticity: The Role of IL-23 in Th17 Fate. J. Cell. Immunol. 4, (2022).

49. Morrison, P. J., Ballantyne, S. J. & Kullberg, M. C. Interleukin-23 and T helper 17-type responses in intestinal inflammation: from cytokines to T-cell plasticity. Immunology 133, 397–408 (2011).

50. Zhao, W. et al. Aberrant methylation of the IL-12B promotor region contributes to the risk of developing ovarian endometriosis. Mol. Reprod. Dev. 86, 632–638 (2019).

51. Mazzeo, D. et al. Interleukin-12 and Its Free p40 Subunit Regulate Immune Recognition of Endometrial Cells: Potential Role in Endometriosis 1. J. Clin. Endocrinol. Metab. 83, 911–916 (1998).

52. Tarokh, M. et al. Serum and Peritoneal Fluid Cytokine Profiles in Infertile Women with Endometriosis. Iran. J. Immunol. 16, 151–162 (2019).

53. Zhang, Q.-F., Chen, G.-Y., Liu, Y., Huang, H.-J. & Song, Y.-F. Relationship between resistin and IL-23 levels in follicular fluid in infertile patients with endometriosis undergoing IVF-ET. Adv. Clin. Exp. Med. 26, 1431–1435 (2017).

54. Andreoli, C. G. et al. T helper (Th)1, Th2, and Th17 interleukin pathways in infertile patients with minimal/mild endometriosis. Fertil. Steril. 95, 2477–2480 (2011).

55. Sugahara, T. et al. Reduced innate lymphoid cells in the endometrium of women with endometriosis. Am. J. Reprod. Immunol. 87, (2022).

56. Montagna, P. et al. Peritoneal fluid macrophages in endometriosis: correlation between the expression of estrogen receptors and inflammation. Fertil. Steril. 90, 156–164 (2008).

57. Symons, L. K. et al. The Immunopathophysiology of Endometriosis. Trends Mol. Med. 24, 748–762 (2018).

58. Harada, T. et al. Increased interleukin-6 levels in peritoneal fluid of infertile patients with active endometriosis. Am. J. Obstet. Gynecol. 176, 593–597 (1997).

59. Naseri, S., Rosenberg-Hasson, Y., Maecker, H. T., Avrutsky, M. I. & Blumenthal, P. D. A cross-sectional study comparing the inflammatory profile of menstrual effluent vs. peripheral blood. Heal. Sci. Reports 6, (2023).

60. Yang, L. et al. IL-21 and TGF-β are required for differentiation of human TH17 cells. Nature 454, 350–352 (2008).

61. McCallion, A. et al. Estrogen mediates inflammatory role of mast cells in endometriosis pathophysiology. Frontiers in Immunology 13, (2022).

62. Lingegowda, H. et al. Implications of dysregulated endogenous cannabinoid family members in the pathophysiology of endometriosis. F&S Sci. 2, 419–430 (2021).

63. Lingegowda, H. et al. Synthetic Cannabinoid Agonist WIN 55212-2 Targets Proliferation, Angiogenesis, and Apoptosis via MAPK/AKT Signaling in Human Endometriotic Cell Lines and a Murine Model of Endometriosis. Front. Reprod. Heal. 3, (2021).

64. Wang, P. et al. IL-23 concentration-dependently regulates T24 cell proliferation, migration and invasion and is associated with prognosis in patients with bladder cancer. Oncol. Rep. (2018). doi:10.3892/or.2018.6775

65. Yan, J., Smyth, M. J. & Teng, M. W. L. Interleukin (IL)-12 and IL-23 and Their Conflicting Roles in Cancer. Cold Spring Harb. Perspect. Biol. 10, a028530 (2018).

66. Suzuki, H. et al. IL-23 directly enhances the proliferative and invasive activities of colorectal carcinoma. Oncol. Lett. 4, 199–204 (2012).

67. Subhadarshani, S., Yusuf, N. & Elmets, C. A. IL-23 and the Tumor Microenvironment. in 89–98 (2021). doi:10.1007/978-3-030-55617-4_6

68. Zhang, W. et al. Transcriptional and posttranslational regulation of Th17/Treg balance in health and disease. Eur. J. Immunol. 51, 2137–2150 (2021).

69. Astry, B., Venkatesha, S. H. & Moudgil, K. D. Involvement of the IL-23/IL-17 axis and the Th17/Treg balance in the pathogenesis and control of autoimmune arthritis. Cytokine 74, 54–61 (2015).

70. Paradowska-Gorycka, A. et al. Th17/Treg-Related Transcriptional Factor Expression and Cytokine Profile in Patients With Rheumatoid Arthritis. Front. Immunol. 11, (2020).

71. Morishima, N., Mizoguchi, I., Takeda, K., Mizuguchi, J. & Yoshimoto, T. TGF-β is necessary for induction of IL-23R and Th17 differentiation by IL-6 and IL-23. Biochem. Biophys. Res. Commun. 386, 105–110 (2009).

72. Ghoreschi, K. et al. Generation of pathogenic TH17 cells in the absence of TGF-β signalling. Nature 467, 967–971 (2010).

73. Tang, C., Chen, S., Qian, H. & Huang, W. Interleukin-23: As a drug target for autoimmune inflammatory diseases. Immunology 135, 112–124 (2012).

74. Yang, K., Oak, A. S. W. & Elewski, B. E. Use of IL-23 Inhibitors for the Treatment of Plaque Psoriasis and Psoriatic Arthritis: A Comprehensive Review. Am. J. Clin. Dermatol. 22, 173–192 (2021).

75. Li, W., Ghamrawi, R., Haidari, W. & Feldman, S. R. Risankizumab for the Treatment of Moderate to Severe Plaque Psoriasis. Ann. Pharmacother. 54, 380–387 (2020).

76. Graier, T. et al. Real-world effectiveness of anti-interleukin-23 antibodies in chronic plaque-type psoriasis of patients from the Austrian Psoriasis Registry (PsoRA). Sci. Rep. 12, 15078 (2022).

77. Yuan, N., Yu, G., Liu, D., Wang, X. & Zhao, L. An emerging role of interleukin-23 in rheumatoid arthritis. Immunopharmacol. Immunotoxicol. 41, 185–191 (2019).

