## Supplementary Figures for "The dysregulated IL-23/T_H_17 axis in endometriosis pathophysiology"

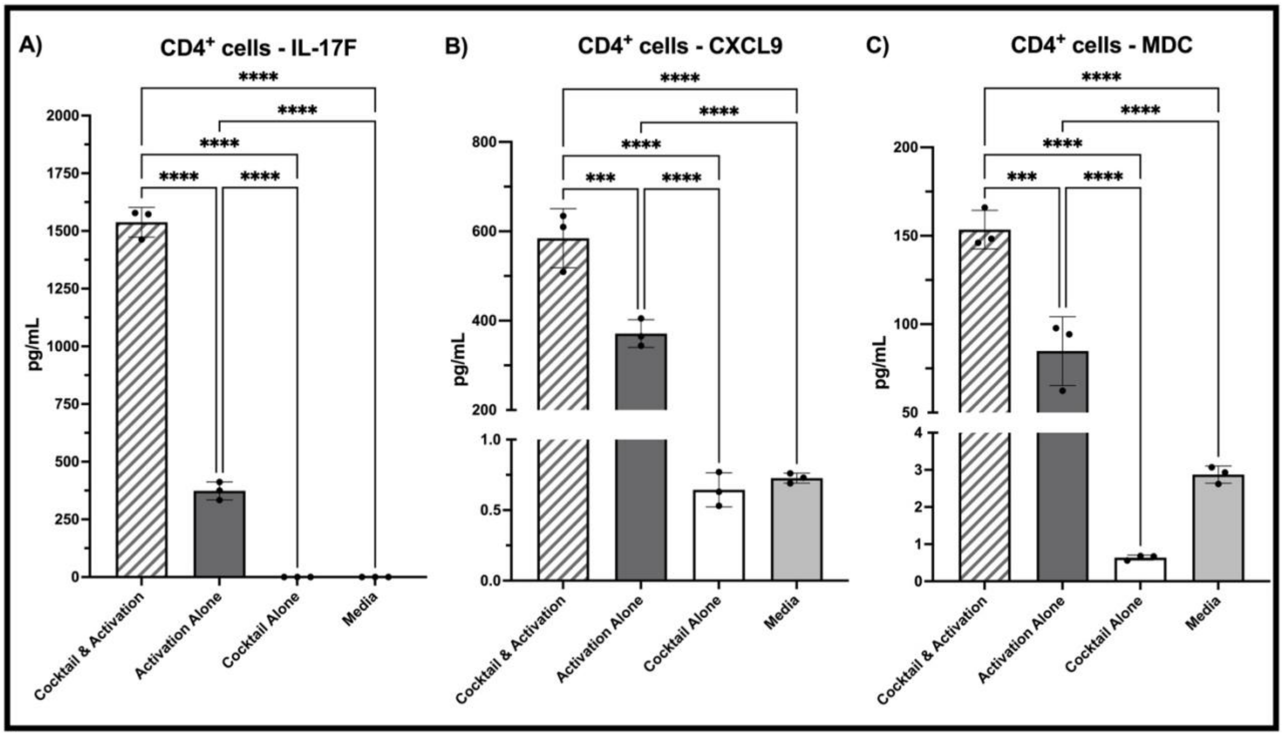


***Supplementary Figure 1:*** ***IL-23 cocktail treatment of primary CD4$^+$ T cells significantly increases the production of factors known to play a role in endometriosis lesion establishment and maintenance.*** *When treated with the IL-23 cocktail (rhIL-23, rhIL-21, rhTGF-β1, rhIL-1β, rhIL-6, anti-IFNγ, and anti-IL-4), primary CD4$^+$ T cells produced significantly increased IL-17F, CXCL9, and MDC, characteristic of pathogenic $T_H$17 cells (**A-C**). Cytokine/chemokine production was detected via multiplex cytokine array. Data is represented as mean ± SD. A one-way ANOVA with Tukey post-hoc was used to assess significance. ***p<0.001, ****p<0.0001.*

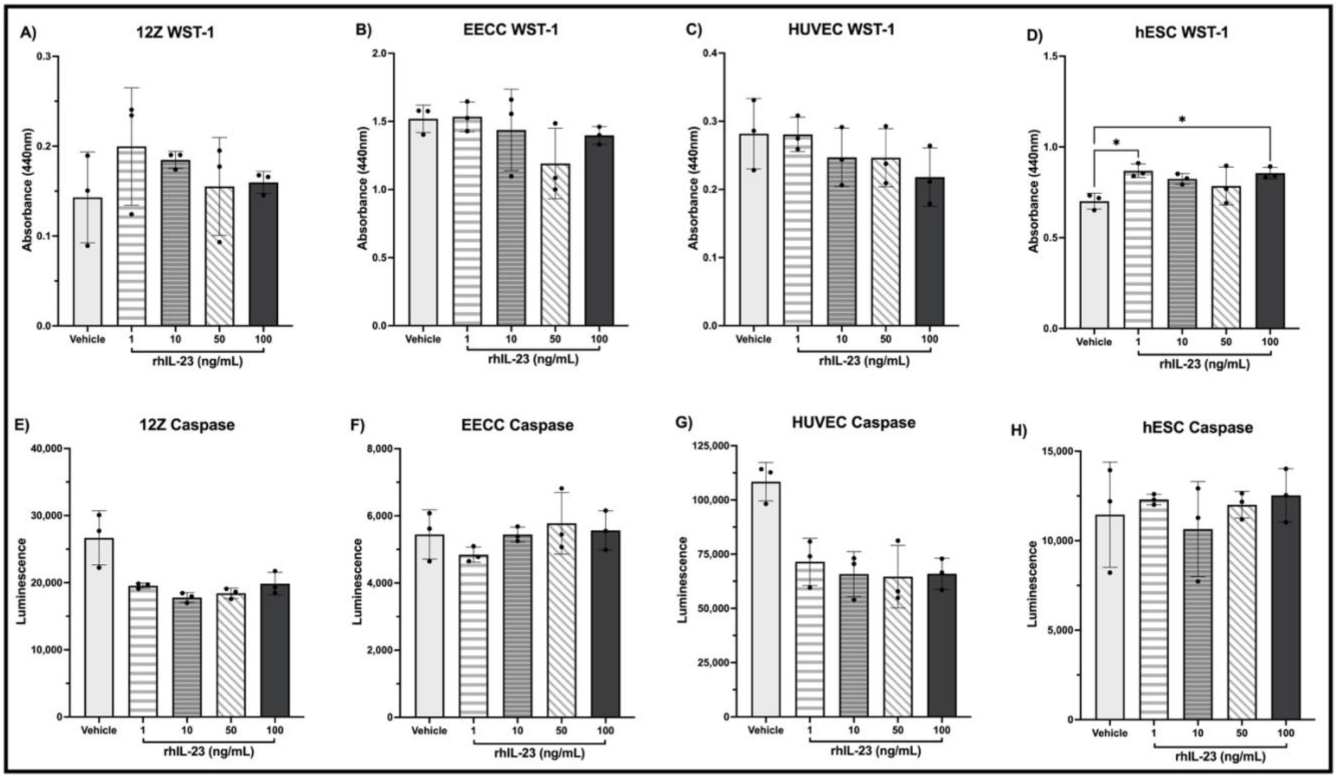


**Supplementary Figure 2:** ***IL-23 treatment significantly increases proliferation of hESCs but did not alter proliferation or apoptosis of other cells representative of the endometriotic microenvironment.*** *Cell lines were treated with 0.01% BSA in PBS (vehicle) or differing concentrations of rhIL-23 (1, 10, 50, 100ng/mL) for 24hrs prior to WST-1 (proliferation) and caspase (apoptosis) assays. Results illustrate that IL-23 treatment does not induce proliferation of 12Z, EECC, or HUVEC cell lines in-vitro as compared to the vehicle group, though IL-23 does significantly increase hESC cell proliferation (****A-D****). Additionally, IL-23 treatment does not significantly alter apoptosis of 12Z, EECC, HUVEC, and hESC cells in-vitro as compared to the vehicle group (****E-H****). A one-way ANOVA with Tukey post-hoc was used to assess significance. *p<0.05.*

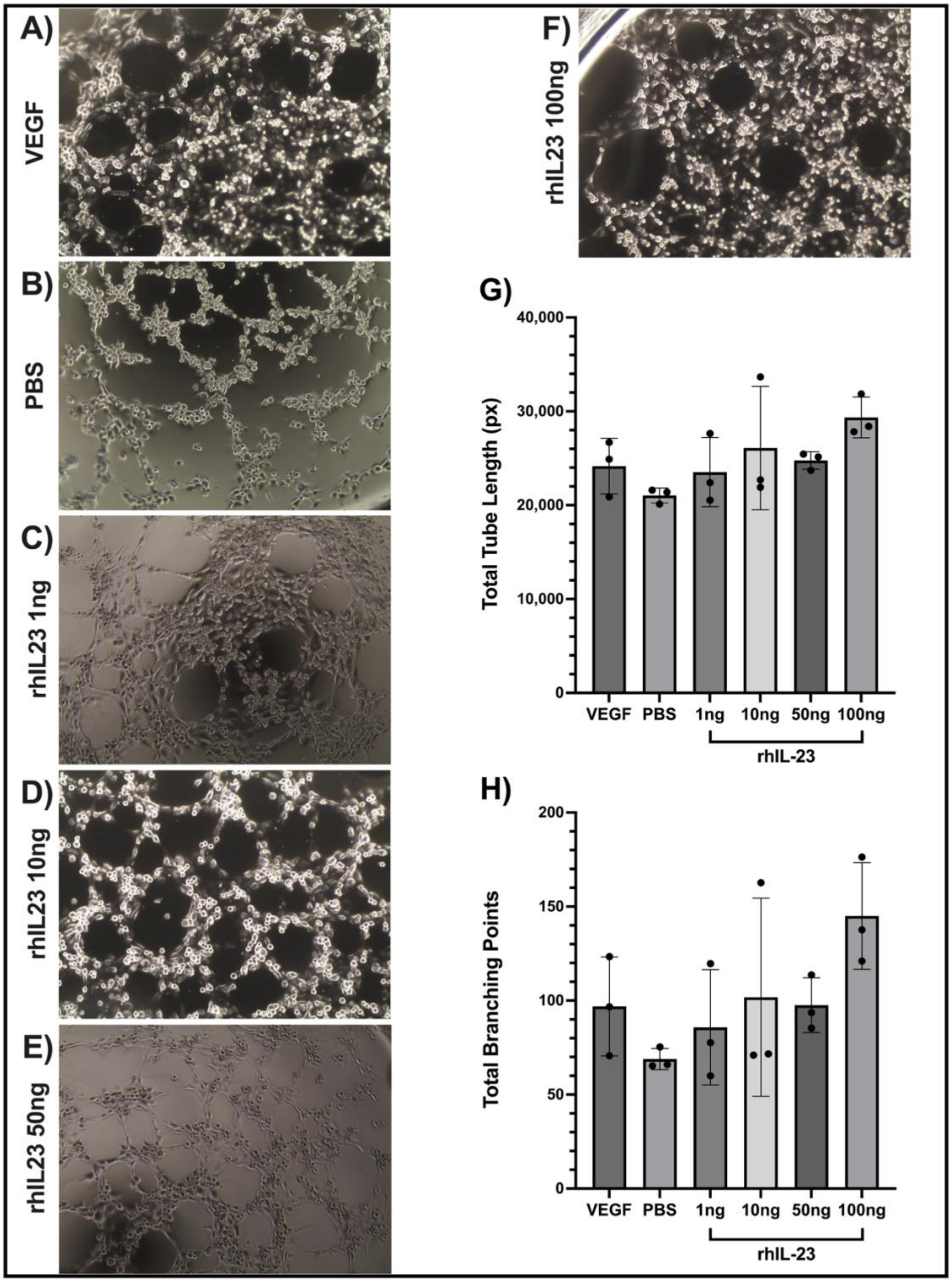


<u>**_Supplementary Figure 3:_**</u> ***IL-23 treatment does not significantly alter tubulogenesis of HUVECs.*** *HUVECs were treated with either VEGF (25ng/mL), PBS, or rhIL-23 (1, 10, 50, 100ng/mL) and incubated for 4hrs prior to image acquisition at 4X objective (**A**-**F**). WIMASIS Software was used to analyze images for total tube length (**G**) and total branching points (**H**). A one-way ANOVA with Tukey post-hoc was used to assess significance.*

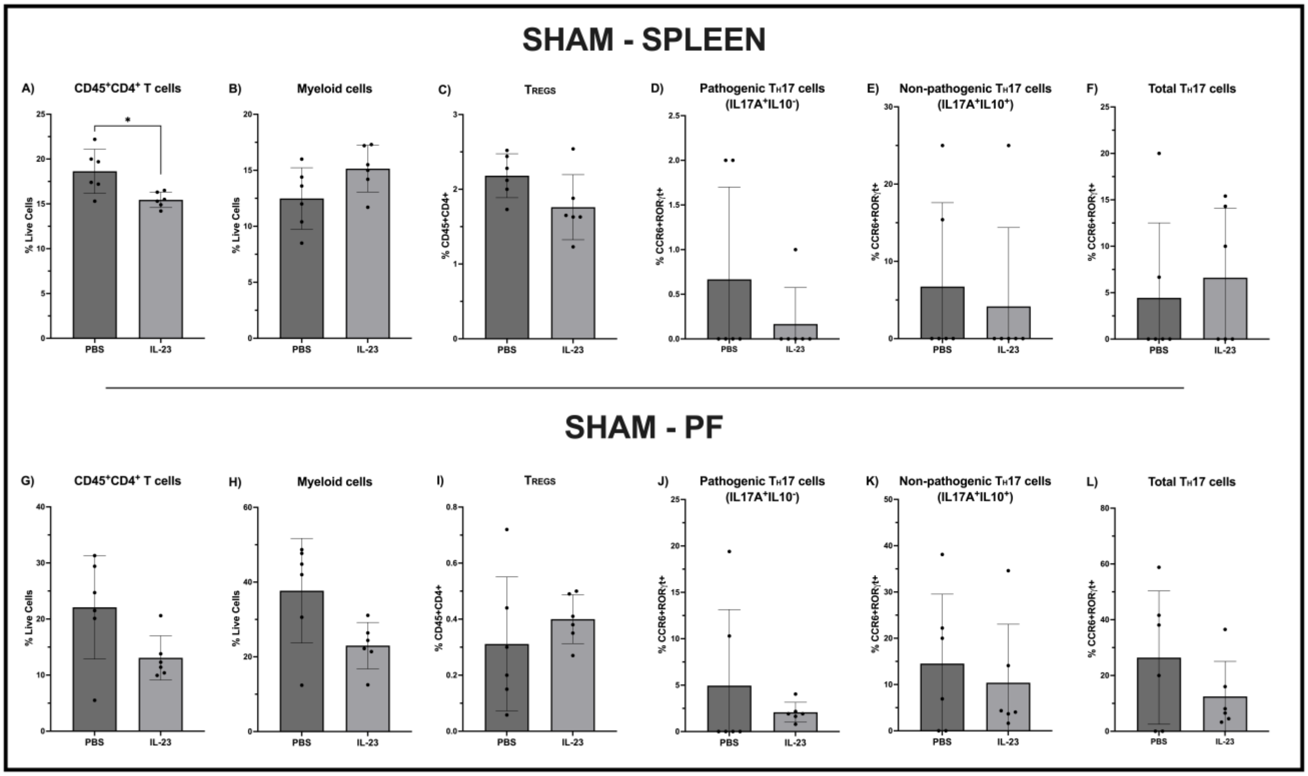


<u>**Supplementary Figure 4:**</u> ***IL-23 treatment results in a significant decrease in CD4$^{+}$ T cells within splenocytes of sham operated mice as compared to controls (PBS), but otherwise does not significantly alter immune cell subsets.*** *Mice (n=12) underwent sham surgery in which no endometrial fragments were adhered to the peritoneal wall. 1 week following surgery, mice were treated (i.p.) with 1µg rmIL-23 (n=6) or PBS (n=6) 3 times a week for 3 weeks before splenocytes and PF were isolated for flow cytometric analysis. Markers were used to detect CD4$^{+}$ T cells (CD45$^{+}$CD4$^{+}$), myeloid cells (CD45$^{+}$CD11b$^{+}$), $T_{REGS}$ (CD45$^{+}$CD4$^{+}$CD25$^{+}$FoxP3$^{+}$), pathogenic $T_{H}$17 cells (CD45$^{+}$CD4$^{+}$CCR6$^{+}$RORγt$^{+}$IL17A$^{+}$IL10$^{-}$), non-pathogenic $T_{H}$17 cells (CD45$^{+}$CD4$^{+}$CCR6$^{+}$RORγt$^{+}$IL17A$^{+}$IL10$^{+}$), and total $T_{H}$17 cells (CD45$^{+}$CD4$^{+}$CCR6$^{+}$RORγt$^{+}$IL17A$^{+}$). Results depict respective immune cell populations within the spleen (**A-F**) as well as the PF (**G-L**). Gating strategy is as depicted in Figure 6. Data is represented as mean ± SD. A non-parametric student's t-test with Mann-Whitney correction was used to assess significance. *$p<0.05$.*

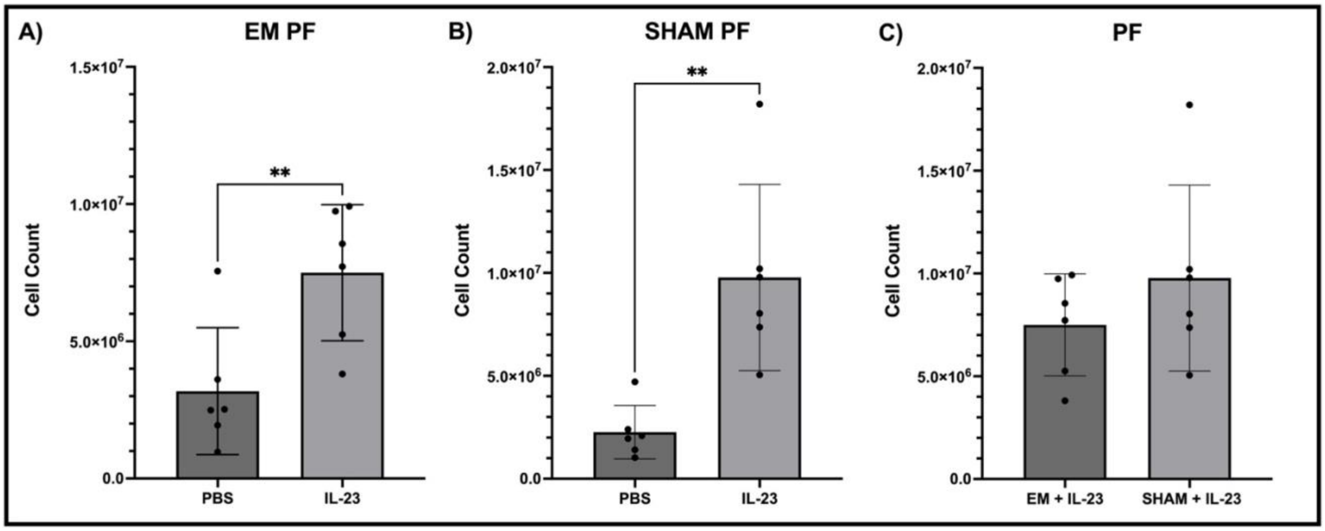


***Supplementary Figure 5:* IL-23 treatment results in significant immune cell recruitment to the peritoneal cavity in both endometriosis-induced and sham operated mice.** *Endometriosis (EM)-induced mice (n=12) were treated (i.p.) with 1µg rmIL-23 (n=6) or PBS (n=6) 3 times a week for 3 weeks before PF collection. Both EM (**A**) and sham (**B**) mouse models had significantly increased PF cells when treated with IL-23 as compared to PBS (controls) after counting PF via automated Countess™ 3 (ThermoFisher, Canada). When comparing EM and sham models treated with IL-23 there were no significant differences found in PF cell counts (**C**). Data is represented as mean ± SD. A non-parametric student's t-test with Mann-Whitney correction was used to assess significance. **p<0.01.*
